# Physiological Substrates and Ontogeny-Specific Expression of the Ubiquitin Ligases MARCH1 and MARCH8

**DOI:** 10.1101/2021.04.15.439921

**Authors:** Patrick Schriek, Haiyin Liu, Alan C. Ching, Pauline Huang, Nishma Gupta, Kayla R. Wilson, MinHsuang Tsai, Yuting Yan, Christophe F. Macri, Laura F. Dagley, Giuseppe Infusini, Andrew I. Webb, Hamish McWilliam, Satoshi Ishido, Justine D. Mintern, Jose A. Villadangos

## Abstract

MARCH1 and MARCH8 are ubiquitin ligases that control the expression and trafficking of critical immunoreceptors. Understanding of their function is hampered by three major knowledge gaps: (i) it is unclear which cell types utilize these ligases; (ii) their level of redundancy is unknown; and (iii) most of their putative substrates have been described in cell lines, often overexpressing MARCH1 or MARCH8, and it is unclear which substrates are regulated by either ligase *in vivo*. Here we address these questions by systematically analyzing the immune cell repertoire of MARCH1- or MARCH8-deficient mice, and applying unbiased proteomic profiling of the plasma membrane of primary cells to identify MARCH1 and MARCH8 substrates. Only CD86 and MHC II were unequivocally identified as immunoreceptors regulated by MARCH1 and MARCH8, but each ligase carried out its function in different tissues. MARCH1 regulated MHC II and CD86 in professional and “atypical” antigen presenting cells of hematopoietic origin, whereas MARCH8 only operated in non-hematopoietic cells. Our results reveal that the range of cells constitutively endowed with antigen-presentation capacity is wider than generally appreciated. They also establish MARCH1 and MARCH8 as specialized regulators of CD4+ T cell immunity in two ontogenically distinct cellular compartments.

## INTRODUCTION

Ubiquitination is a major mechanism for the regulation of membrane proteostasis. In brief, covalent attachment of ubiquitin (Ub) chains to the cytosolic tail of transmembrane proteins promotes endosomal trafficking to multivesicular bodies for subsequent degradation in lysosomes [1]. This post-translational modification enables the fine-tuning of surface protein expression levels. Ub is attached to substrates by E3 Ub ligases. Membrane Associated RING-CH Finger (MARCH, gene symbol *Marchf*) is a family of eleven E3 ligases, all of which possess two or more transmembrane domains, with the exception of MARCH7 and MARCH10 [2]. They were initially identified as the mammalian homologues of herpesvirus immunoevasins that ubiquitinate host molecules involved in anti-viral immunity to subvert immune responses [3][4]. MARCH E3 Ub ligases are thought to be specialized at ubiquitinating immunoregulatory receptors, but their physiological substrates remain largely unknown [2][5]. It is also unclear if their expression and function is restricted to cells of the immune system and, if so, which.

MARCH1 and MARCH8 are the most studied members of the MARCH family. As they share approximately 60% overall sequence homology [2], they are thought to also share substrate specificity. Indeed, both ubiquitinate major histocompatibility complex class II (MHC II) molecules, the receptor employed by antigen presenting cells (APC) to display peptide antigens to CD4^+^ T cells. By regulating MHC II expression [6][7][8], MARCH1 and MARCH8 play key roles in CD4^+^ T cell development in the thymus [9][10][11] and activation in the periphery [12], respectively. Furthermore, they have been involved in complex immune reactions such as inflammation [13], immunity to infection [14][15], cancer [16], allergy and autoimmunity [17][18]. This poses the question whether both ligases regulate the expression of other immune receptors, some of which reportedly include CD44 [19], CD71 [20], CD86 [21], CD95 [22] and CD98 [19] among others [5][23]. However, to date CD86 is the only membrane protein apart from MHC II that has been shown to be regulated by MARCH1 *in vivo* [21], and it is not known if it can also be regulated by MARCH8. All other putative MARCH1 or MARCH8 substrates have been described in cell lines and/or overexpression studies. MARCH proteins are expressed at very low levels in primary cells [2][24][25][26], and since E3 ligase overexpression can cause off-target effects, it remains unclear which, if any of the MARCH1 and MARCH8 substrates described in transfected cell lines are ubiquitinated by these ligases in physiological settings. To summarize, the repertoire of MARCH1 and MARCH8 substrates *in vivo* remains largely unknown. This is an important shortcoming because ubiquitination is amenable to pharmacological manipulation [27][28], and development of drugs targeting MARCH1 or MARCH8 might have therapeutic potential provided their substrates are identified.

Another important knowledge gap in MARCH1 and MARCH8 biology pertains to their expression pattern. Quantitating MARCH1 or MARCH8 protein expression is unfeasible due to their low abundance [2] and fast turn-over [29][30], and even their transcription levels are poor predictors of function [24][25][26]. Identification of MARCH1- or MARCH8-expressing cells thus relies on analysis of surface expression of membrane protein substrates as a surrogate of activity. MARCH1 ubiquitinates MHC II and CD86 in B cells and conventional and plasmacytoid dendritic cells (cDC and pDC, respectively) [6][7][8], but it is not functional in thymic epithelial cells (TEC) [9][10]. Whether it is active in other hematopoietic or non- hematopoietic cells remains unknown. In contrast, MARCH8 ubiquitinates MHC II in TEC, not in B cells or DC [9][10], but it is not known if it ubiquitinates other receptors in these cells, and whether it is also expressed in other cells. Incomplete understanding of the pattern of MARCH expression again limits the development and potential application of ubiquitination- modulating agents as immunomodulatory drugs.

Here, we present a systematic analysis of the pattern of activity of MARCH1 and MARCH8 in multiple hematopoietic and non-hematopoietic cells isolated from *Marchf1*^−/−^ and *Marchf8*^−/−^ mice. We have also carried out quantitative proteomic comparisons of WT vs *Marchf1*^−/−^ or *Marchf8*^−/−^ plasma membrane purified from cDC and B cells. Our results define physiological substrates regulated by these two ligases and demonstrate functional specializations of MARCH1 and MARCH8 in two ontogenically distinct compartments.

## MATERIALS AND METHODS

### Mice

Wild type (WT, C57BL/6), *Marchf1*^−/−^ [31], *Marchf8*^−/−^ [9] and *I-Aα*^−/−^ [32] mice were bred and maintained in specific pathogen-free conditions within the Melbourne Bioresources Platform at the Bio21 Molecular Science and Biotechnology Institute. Analyses were undertaken with male or female mice aged between 6-14 weeks and performed in accordance with the Institutional Animal Care and Use Committee guidelines of the University of Melbourne. All procedures were approved by the Animal Ethics Committee at the University of Melbourne.

### Isolation of mouse primary cells and analytical flow cytometry

Single cell suspensions from blood, spleen, subcutaneous lymph nodes (LN), thymus, peritoneal cavity and lung were generated for analysis of B cells, T cells, DC, granulocytes, macrophages, monocytes, neutrophils, eosinophils and thymic or alveolar epithelial cells. Blood was collected from submandibular veins and red blood cells were lysed. Whole single cell suspensions from spleen and subcutaneous LN (axillary and inguinal) were generated by spleen digestion with 0.1 % DNase I (Roche) and 1 mg/ml collagenase type III (Worthington) and red blood cell lysis. DCs from spleen and LN were further enriched by selection of low-density cells by density gradient centrifugation in 1.077 g/cm^3^ Nycodenz® (Axis shield). Thymi were digested in 0.1 % DNase I (Roche) and 0.5 U/ml liberase (Roche) and thymic cDC were further enriched by 1.077 g/cm^3^ Nycodenz® density gradient centrifugation (Axis shield). Cells from the peritoneal cavity were harvested by injection and aspiration of PBS. Lungs were perfused with PBS and digested with 50 µg/ml DNase I (Roche) and 0.25 mg/ml liberase (Roche) and red blood cells lysed.

For flow cytometry, cells were incubated with FcR blocking reagent (Miltenyi Biotec), prior to staining with mAb detecting B220/CD45R (RA3-6B2), CD19 (6D5), CD64 (X54-5/7.1), F4/80 (F4/80, Walter Eliza Hall Institute (WEHI) Antibody Facility), CD3 (KT3-1.1, WEHI Antibody Facility), TCRβ (Η57-597, WEHI Antibody Facility), CD4 (GK1.5), CD8 (YTS169.4 WEHI Antibody Facility), CD8 (53-6.7), BST-2 (927), Siglec-H (551), MHC II (M5/114), CD11c (N418), CD11b (M1/70), Ly6G (1A8), Ly6C (HK1.4), NK1.1 (PK136, BD Biosciences), Sirpα (P84), XCR1 (ZET), CD45 (30-F11), EpCAM (G8.8), Ly51 (6C3), UEA-1 (Vector Laboratories), MerTK (2B10C42), Siglec-F (E50-2440 BD Biosciences), CD31 (390), CD24 (M1/69, WEHI Antibody Facility), Sca-1 (D7), CD86 (GL-1), CD40 (FGK45.5, Miltenyi Biotec), CD80 (16-10A1, BD Biosciences), CD44 (IM7.81), CD71 (R17217, eBiosciences), CD95 (15A7, eBiosciences), CD98 (RL388), PD-L1 (10F.9G2), PD-L2 (TY25), ICOS-L (HK5.3), B7-H3 (MIH35) or B7-H4 (HMH4-5G1), conjugated to fluorochromes BUV395, BUV805, FITC, PE, PE-Cy7, PerCP/Cy5.5, APC, APC-Cy7, AF700, BV785, BV650, BV510 or BV421 (all from BioLegend, if not stated differently). Cell viability was determined with Fixable Viability Dye eFluor™ 780 (eBiosciences), propidium iodide (PI) or diamidino phenylindole (DAPI). Analysis was performed using a LSRFortessa (BD Biosciences) or CytoFLEX LX (Beckman Coulter) in the Melbourne Cytometry Platform (University of Melbourne). Data was analyzed with FlowJo (Tree Star) and GraphPad Prism. **Supplementary Figures 1 and 2** summarize gating strategies for cells from blood, spleen, subcutaneous lymph nodes (LN), thymus, peritoneal cavity and lung.

### Isolation of primary immune cells for proteomic analysis

B cells were purified from spleens using Ficoll® Paque Plus (GE Healthcare) gradient centrifugation and negative depletion with FITC-conjugated mAb specific for CD4 (GK1.5), Ly-76 (TER119) and CD43 (S7) and magnetic anti-FITC MicroBeads (Miltenyi Biotec). Preparations were approximately 95-98% pure for CD19^+^ B220^+^ B cells. Splenic cDC were purified from mice subcutaneously injected with Flt3L-secreting melanoma cells [33], 9 days before purification. cDC were purified from spleens of Flt3L-expanded mice following spleen digestion with DNase I (Roche) and collagenase type III and Nycodenz® density gradient centrifugation (Axis shield) with subsequent negative depletion using rat mAb specific for CD3 (KT3-1.1), Thy1 (T24/31.7), Ly-76 (Ter119), B220 (RA3-6B2) and Ly-6C/G (RB6-8C5) and anti-rat IgG-coupled magnetic beads (Qiagen) as previously described [34]. Preparations were approximately 90-95% pure for CD11c^+^ MHC II^+^ cDC.

### Preparation of subcellular fractions enriched in plasma membrane and intracellular compartments for proteomics

Subcellular fractionation was performed as previously described [35]. In brief, purified B cells (4-5 x 10^7^ cells, 95-98% purity) and cDC (4-5 x 10^7^ cells, 90-95% purity) from spleens of WT, *Marchf1*^−/−^ and *Marchf8*^−/−^ mice were incubated with FITC-conjugated anti-CD19 and anti-B220 mAb (B cells) or anti-CD11c, anti-CD45.2, anti-CD49d and anti-MHC I mAb (cDC). mAb- labelled cells were homogenized in the presence of cOmplete^TM^ protease inhibitors (Roche) by mechanical disruption using a cell-cracker (HGM Laboratory equipment). Homogenized preparations were centrifuged at low speed to obtain post-nuclear supernatant (PNS). Surface-labelled plasma membrane (PM) microsomes were isolated by magnetic immunoaffinity using anti-FITC mAb-coated magnetic beads (Miltenyi Biotec) and concentrated by ultracentrifugation in thickwall polycarbonate tubes (Beckman Coulter). PNS with the PM fraction removed was likewise ultracentrifuged to sediment the “intracellular compartments” (IC) fraction.

### Proteomic profiling of differentially expressed PM proteins

Subcellular fractions (PM and IC) were prepared for mass spectrometry analysis from three independent cell preparations using FASP protein digestion (Protein Discovery) as previously described [36], with the following modifications. Proteins were reduced and digested with sequence-grade modified Trypsin Gold (Promega). Peptides were eluted with ammonium bicarbonate and acidified peptide mixtures from each biological replicate were analyzed in technical triplicates by nanoflow reverse-phase liquid chromatography tandem mass spectrometry (LC-MS/MS) on a nanoAcquity system (Waters) coupled to a Q-Exactive mass spectrometer equipped with a nanoelectrospray ion source for automated MS/MS (Thermo Fisher Scientific). High-resolution MS/MS spectra were processed with MaxQuant (version 1.6.7.0) for feature detection and protein identification using the Andromeda search engine [37]. Extracted peak lists were searched against the UniProtKB/Swiss-Prot Mus musculus database (Oct-2019) and a separate reverse decoy database to empirically assess the false discovery rate (FDR) using a strict trypsin specificity allowing up to 2 missed cleavages. The minimum required peptide length was 7 amino acids. The “match between runs” option in MaxQuant was used [38]. PSM and protein identifications were filtered using a target-decoy approach at a FDR of 1%. LFQ quantification was performed, with a minimum ratio of 2. Protein relative quantitative analysis was performed in R using MaxQuant’s proteinGroups.txt and LFQ intensities. Missing values were imputed using a random normal distribution of values derived from the measured distribution of intensities [39] using a mean with a negative shift of 1.8 standard deviations and a standard deviation equal to 0.3 of the standard deviation of the measured intensities. The probability of differential expression was calculated using the function *lmFit* from the Bioconductor package limma [40] followed by *eBayes* using the default settings [41] and false-discovery rate correction using the Benjamini–Hochberg method. The output included P value, confidence interval and ratio estimate. GO-term enrichment analysis was performed using the *enrichr* function in the Bioconductor clusterProfiler package [42]. Enrichment was calculated separately for the proteins overrepresented in each fraction, relative to all proteins identified in collected fractions across all the LCMS runs, and GO term association was filtered to include only experimental and high throughput evidence. Enrichment P values were corrected for multiple testing using the function’s *‘fdr’* method. The mass spectrometry proteomics data have been deposited to the ProteomeXchange Consortium via the PRIDE [43]. The PRIDE database and related tools and resources in 2019: improving support for quantification data. Nucleic Acids Res 47(D1):D442- D450 partner repository with the dataset identifier *PXD023115*.

## RESULTS

### MARCH1, but not MARCH8, is functional in professional APC

The first objective of this study was to establish which mouse cells express MARCH1 or MARCH8. Their low level of transcription combined with fast turn-over contribute to maintain the two proteins at non-detectable levels in primary cells, hampering definition of their expression pattern. We reasoned that MHC II and/or CD86 could be used as reporters of MARCH1 and MARCH8 activity because in all primary or transformed cells analyzed so far, the surface level of these two receptors decreases by expression of either ligase [44]. Cells that express MHC II or CD86 and either MARCH1 or MARCH8 should therefore display higher levels of the receptor(s) in *Marchf1*^−/−^ or *Marchf8*^−/−^ mice.

First, we examined professional APC (defined as cells that express detectable levels of MHC II in the steady-state [45][46]) and T cells across various tissues. B cells, cDC1, cDC2, pDC and macrophages from blood, spleen, subcutaneous lymph nodes (LN), thymus, peritoneal cavity and lung of *Marchf1*^−/−^ mice displayed elevated surface MHC II and CD86 relative to WT cells, while no changes were observed in their *Marchf8*^−/−^ counterparts (**Figure 1A-F**). CD4^+^ and CD8^+^ T cells in spleen and LN showed no detectable surface MHC II and their CD86 expression [47] was not altered by MARCH1- nor MARCH8-deficiency (**Figure 1B-C**). MHC II and CD86 expression in peritoneal cDC deficient in both MARCH1 and MARCH8 (*Marchf1*^−/−^ x *Marchf8*^−/−^) was not elevated above that of *Marchf1*^−/−^ cells (**Supplementary Figure 3)**. These results indicate that MARCH1 is expressed and active in all professional APC across various organs/tissues whereas MARCH8 is not or, if it is, does not display enough activity to compensate for the loss of MARCH1.

**Figure 1.**
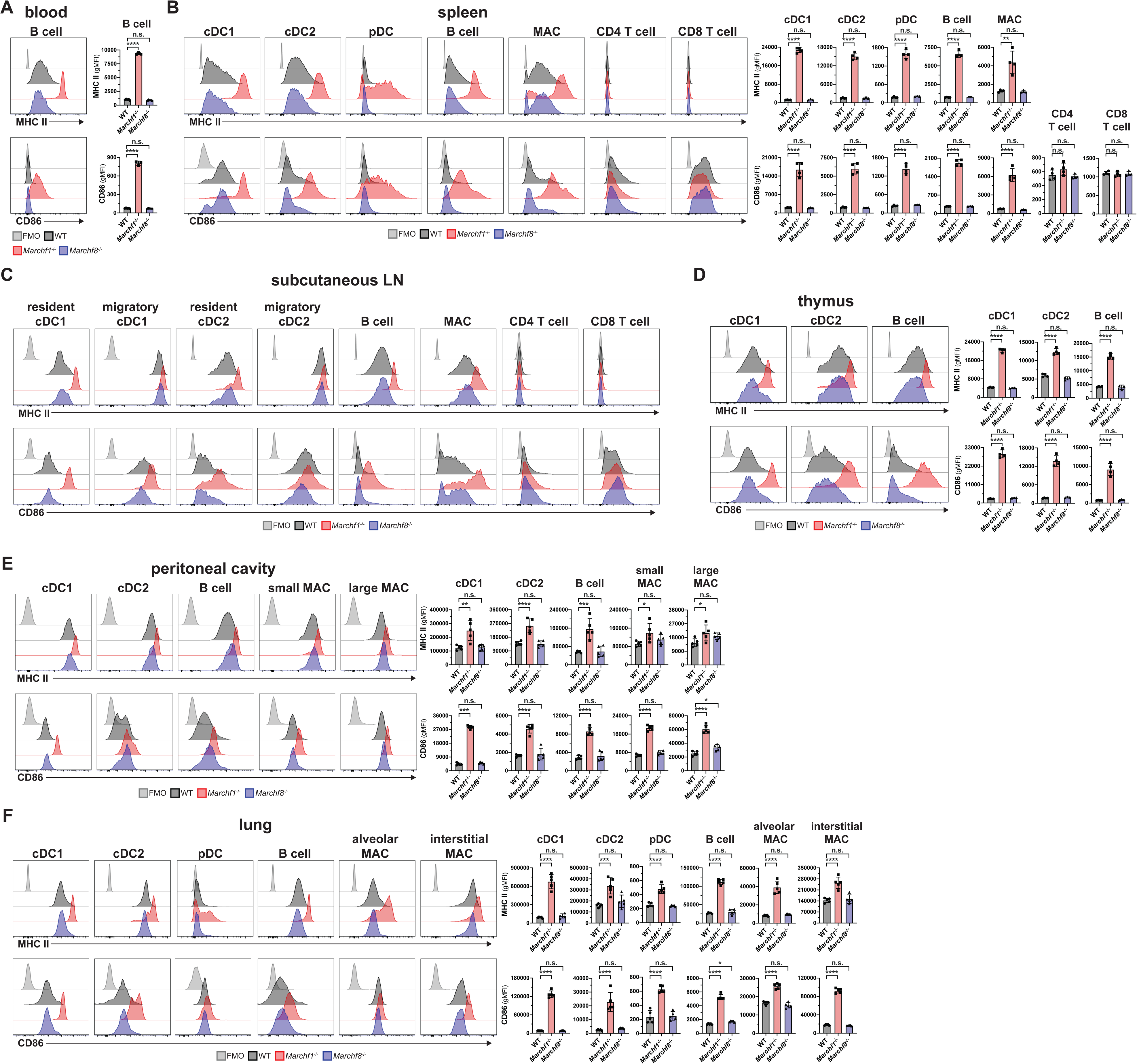
Ubiquitination of MHC II and CD86 by MARCH1 and MARCH8 in haemopoietic professional antigen presenting cells. Surface expression of MHC II and CD86 in (A) blood B cells, (B) splenic cDC1, cDC2, pDC, B cells, macrophages (MAC) and CD4^+^/CD8^+^ T cells, (C) resident and migratory cDC1 and cDC2 as well as B cells, macrophages and CD4^+^/CD8^+^ T cells in subcutaneous (axillary + inguinal) lymph nodes, (D) thymic cDC1, cDC2 and B cells, (D) peritoneal cDC1, cDC2, B cells and small/large macrophages and (E) lung cDC1, cDC2, pDC, B cells and alveolar/interstitial macrophages, all purified from WT mice or mice deficient in either MARCH1 or MARCH8. In all cases a fluorescence-minus-one (FMO) control was included, for which cells were incubated with the corresponding multi-colour staining panel, excluding the fluorescently labelled antibody species of interest (i.e. anti-CD86 or anti-MHC II mAb). Bars represent mean ± SD with each symbol representing an individual mouse (n=4-5). Statistical analysis was performed using one-way ANOVA followed by Sidak’s multiple comparisons test. **** p < 0.0001, *** p < 0.0002, ** p < 0.002, * p < 0.03, n.s. not significant.

Next, we assessed the contribution of MARCH1 to activation-dependent regulation of MHC II and CD86 expression in cDC, the archetypical professional APC. Toll-like receptor (TLR) ligands trigger an activation program in DC, known as DC maturation, that includes up- regulation of MHC II and CD86 expression on the plasma membrane, among other receptors [48]. Activation also leads to down-regulation of *Marchf1* transcription which, combined with fast turn-over of MARCH1, results in negligible expression of the protein in activated DCs [6][49][50][51]. It has been assumed that this change is responsible for the accumulation of MHC II and CD86 on the plasma membrane during cDC activation, but this has not been directly examined. If ubiquitination were the dominant mechanism controlling how much MHC II and CD86 is displayed on cDC, it would be expected that the expression of these two molecules would not vary during activation of *Marchf1*^−/−^ cDC. However, activation of *Marchf1*^−/−^ cDC further increased surface expression of MHC II by ∼1.5 times, and increased CD86 by ∼4 times, when compared than their resting counterparts (**Figure 2**). CD40, which also increases in expression during activation, though it is not a MARCH1 substrate, was expressed at equivalent levels in WT and *Marchf1*^−/−^ cDC at both resting and activated states, so up-regulation of MHC II and CD86 in *Marchf1*^−/−^ cDC could not be attributed to overall dysregulation of surface receptor expression (**Figure 2**). These results indicate that the main contributor to MHC II and, especially, CD86 up-regulation during DC activation is not reduced ubiquitination and degradation, but sustained deposition of newly synthesized molecules on the cell surface [52][53]. DC lacking MARCH8 were indistinguishable from WT cDC in these experiments, again indicating it has no role in resting or activated cDC (**Figure 2**).

**Figure 2.**
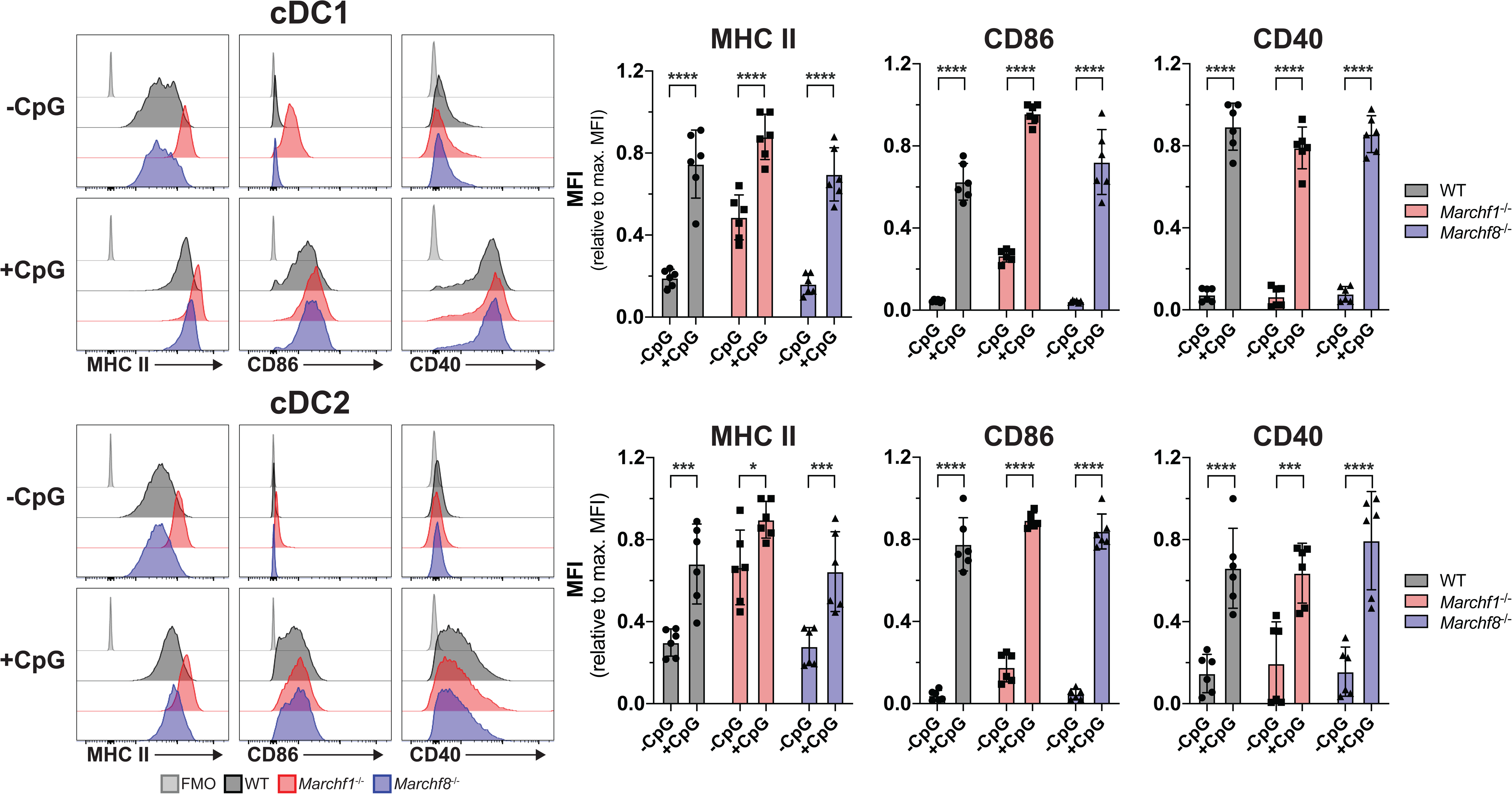
The role of ubiquitination of MHC II and CD86 by MARCH1 and MARCH8 in DC maturation. (A) Surface expression of MHC II and CD86 in CpG-activated cDC purified from the spleen of WT mice or mice deficient in either MARCH1 or MARCH8. Purified splenic cDC (2×10^5^ cells) were incubated for 16 hours *ex vivo* with or without 50 nm CpG in 96-well plates, then washed and analyzed by flow cytometry for MHC II and CD86 surface expression. A fluorescence-minus-one (FMO) control was included, for which cells were incubated with the corresponding multi-colour staining panel, excluding the fluorescently labelled antibody species of interest (i.e. anti-CD86 or anti-MHC II mAb). Bars represent mean ± SD with each symbol representing an individual mouse (n=6). Statistical analysis was performed using one-way ANOVA followed by Sidak’s multiple comparisons test. **** p < 0.0001, *** p < 0.0002, ** p < 0.002, * p < 0.03, n.s. not significant.

### Granulocytes and monocytes express MHC II and CD86, but MARCH1 ubiquitination maintains their surface expression at negligible levels

Next we assessed MARCH1 and MARCH8 activity in “atypical APC”, this is, immune cells that are not considered professional APC but have been suggested to play antigen-presenting roles under certain conditions [46]. These include neutrophils, eosinophils and “inflammatory” (Ly6C^+^) and “patrolling” (Ly6C^−^) monocytes. While monocytes have the potential to develop into macrophages or DCs in inflamed sites [54], they are not thought to perform antigen presenting functions in their undifferentiated state [55]. We examined these atypical APC in spleen and lung. MHC II expression in WT neutrophils, eosinophils and monocytes was barely detectable by flow cytometry, staining at just above the background level observed in cells of mice that do not express any surface MHC II at all (**Figure 3**). Strikingly, all four cell types deficient in MARCH1 expressed MHC II at levels comparable to WT B cells or cDC (compare **Figures 1B** and **F** to **Figures 3A** and **B**, respectively), though expression was higher in spleen than it was in their lung counterparts (**Figure 3A** and **B)**. CD86 was also highly expressed on all four MARCH1-deficient cell types, in this case both in spleen and lungs (**Figure 3**). MARCH8-deficient cells did not display altered MHC II or CD86 expression, confirming this member of the MARCH family is not expressed and/or active in hematopoietic cells (**Figure 3**). Of note, MARCH1-deficient T cells lacked surface MHC II and did not exhibit enriched CD86 expression when deficient in MARCH1 (**Figure 1B**), so neither mutation caused ectopic or increased expression of either molecule. We conclude that neutrophils, eosinophils, monocytes and possibly other atypical APC types [46] produce receptors for antigen presentation and T cell stimulation constitutively. While MARCH1 ubiquitination maintains the surface expression of these proteins at barely detectable levels, these atypical APC might be capable of CD4^+^ T cell priming under certain conditions.

**Figure 3.**
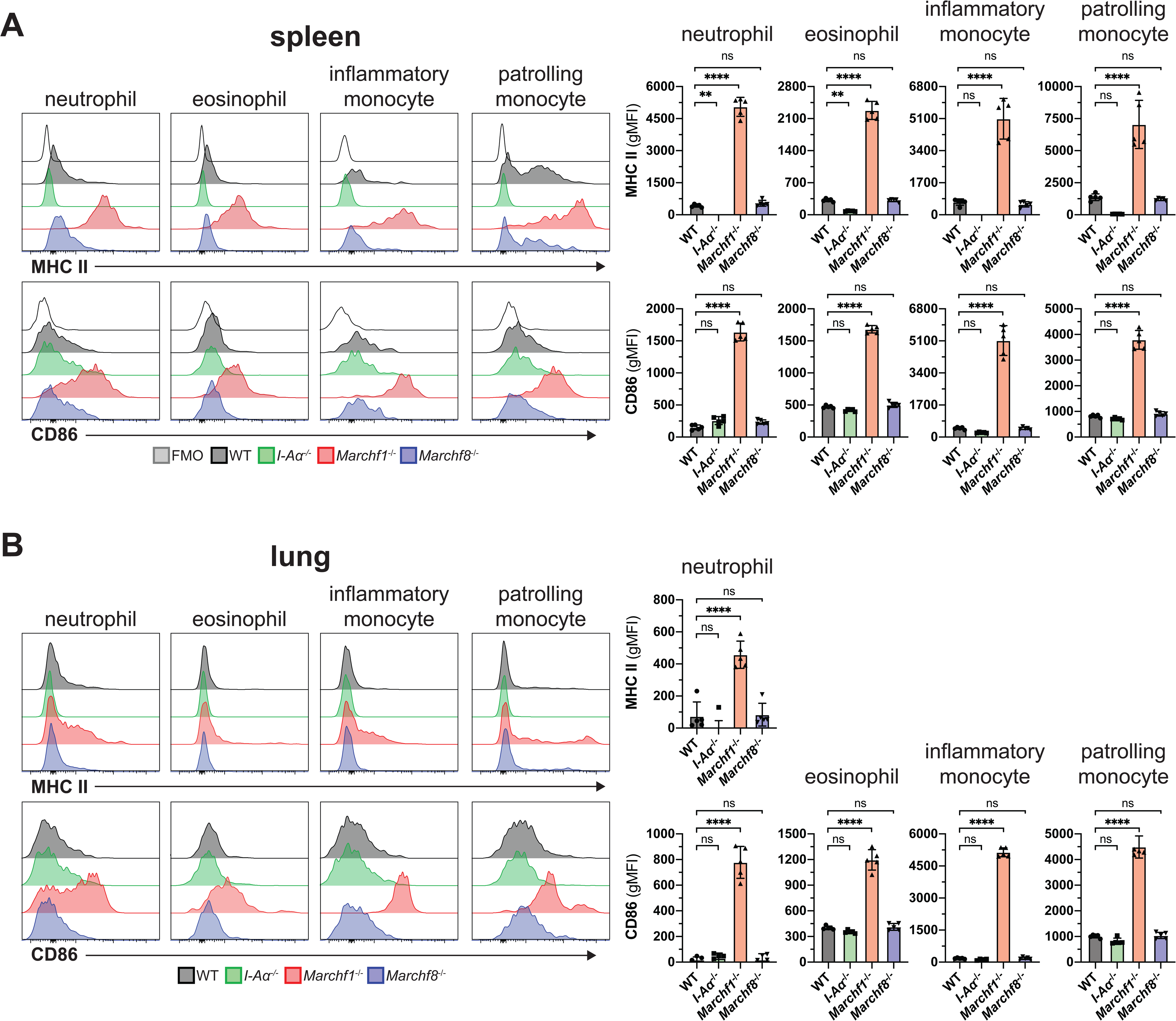
Ubiquitination of MHC II and CD86 by MARCH1 and MARCH8 in granulocytes and monocytes. Surface expression of MHC II and CD86 in neutrophils, eosinophils and inflammatory and patrolling monocytes purified from (A) spleen and (B) lung of WT mice or mice deficient in I-Aα, MARCH1 or MARCH8. Bars represent mean ± SD with each symbol representing an individual mouse (n=5). Statistical analysis was performed using one-way ANOVA followed by Sidak’s multiple comparisons test. **** p < 0.0001, *** p < 0.0002, ** p < 0.002, * p < 0.03, n.s. not significant.

### Previously predicted MARCH1 substrates display normal expression in *Marchf1^−/−^* mice

The second objective of this study was to identify which of the receptors found to be ubiquitinated by MARCH1 or MARCH8 in (transfected) cell lines are also substrates *in vivo* under physiological conditions. Such receptors include CD44, CD71, CD95 and CD98 (reviewed in [5][56]). Carrying out this analysis also allowed us to address the possibility that, contrary to our conclusions above, MARCH8 might be expressed and active in these cells but dedicated to ubiquitinate these receptors rather than MHC II and CD86. This was not the case; expression of CD44, CD71, and CD98 was unaltered in *Marchf8*^−/−^ cDC and B cells compared to WT cells (**Figure 4A**). Furthermore, *Marchf1*^−/−^ cDC and B cells also expressed normal levels of the three receptors (**Figure 4A**). We extended our analysis to other regulatory receptors of T cell activation, including CD40 and members of the B7 family to which CD86 (B7.2) belongs: CD80 (B7.1), CD274 (PD-L1), CD273 (PD-L2), CD275 (ICOS-L), CD276 (B7-H3) and B7-H4. Expression of all these receptors on cDC1, cDC2, pDC and B cells was unaltered in the absence of MARCH1 (**Figure 4B**).

**Figure 4.**
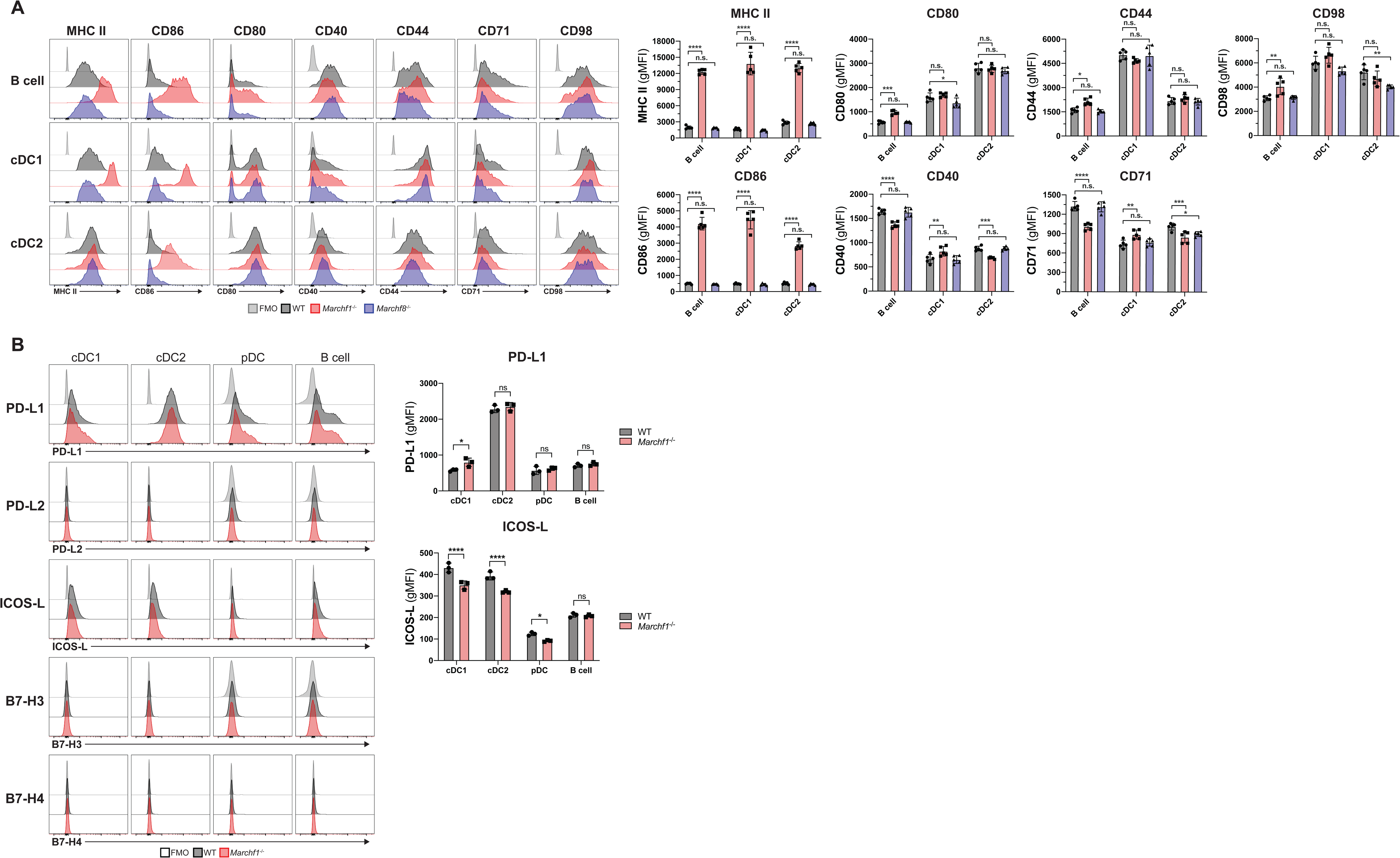
Analysis of putative MARCH1 and MARCH8 substrates in haemopoietic antigen presenting cells. (A) Surface expression of MHC II, CD86, CD80, CD40, CD44, CD71 and CD98 in splenic B cells, cDC1 and cDC2 from WT, *Marchf1*^−/−^ and *Marchf8^−/−^* mice. (B) Surface expression of B7 costimulatory molecules, PD-L1, PD-L2, ICOS-L, B7-H3 and B7-H4, in splenic cDC1, cDC2, pDC and B cells purified from WT or *Marchf1*^−/−^ mice. In all cases a fluorescence-minus-one (FMO) control was included, for which cells were incubated with the corresponding multi-colour staining panel, excluding the fluorescently labelled antibody species of interest. Bars represent mean ± SD with each symbol representing an individual mouse (n=3-6). Statistical analysis was performed using one-way ANOVA followed by Sidak’s multiple comparisons test. **** p < 0.0001, *** p < 0.0002, ** p < 0.002, * p < 0.03, n.s. not significant.

### Proteomic profiling of the plasma membrane of MARCH1- and MARCH8-deficient cDC and B cells

To more comprehensively address the role of MARCH1 and MARCH8 in APC membrane proteostasis, we performed an unbiased proteomic screen where we compared the proteomes of subcellular microsomal fractions enriched in plasma membrane (PM) of WT versus *Marchf1*^−/−^ or *Marchf8*^−/−^ cDC and B cells. We have previously shown this is a robust approach to identify differentially expressed PM proteins between closely related cell populations such as the two major cDC subtypes, cDC1 and cDC2 [57]. To obtain sufficient numbers of primary cDC for this purpose, these cells were expanded in WT, *Marchf1*^−/−^ and *Marchf8*^−/−^ mice bearing a melanoma cell line that secretes the DC growth factor, Flt3L [33]. The cDC expanded using this approach are phenotypically and functionally equivalent to their counterparts in untreated mice [57]. Splenic B cells were purified from untreated mice. The protein profiles of each fraction were identified by semi-quantitative mass spectrometry from three biological replicates, each measured in technical triplicates.

We identified 1868-3108 proteins in the PM fraction of each cell type (**Supplementary Table 1**, total number of IDed proteins regardless of any restrictions). Of note, the subcellular fractions are comprised of microsomes generated during mechanical homogenization of cells, so their composition includes PM but also cytosolic and extracellular content ‘trapped’ inside microsomes or tethered to the cell surface. This method enables analysis of proteins loosely associated with the inner or outer leaflet of the PM. To test the efficiency of the PM-enrichment method, we also sedimented and analyzed in parallel the compartments that remained in the post-nuclear supernatant (PNS) of homogenized cells after retrieval of the PM fraction (mitochondria, endosomes, etc, henceforth termed intracellular compartments, IC). We identified 2073-3537 proteins in the IC fraction of each cell type (**Supplementary Table 2**, total number of IDed proteins regardless of any restrictions). In order to assess enrichment of the PM by this methodology, we compared Gene Ontology (GO) terms/annotations of the proteins identified in the PM and IC fractions of each cell type. This comparison clearly demonstrated enrichment of proteins known to be expressed at the cell surface in the PM fractions, and enrichment of proteins known to occur in intracellular compartments in the IC fractions, validating the subcellular fractionation protocol (**Figure 5A** and **Supplementary Figure 4)**.

**Figure 5.**
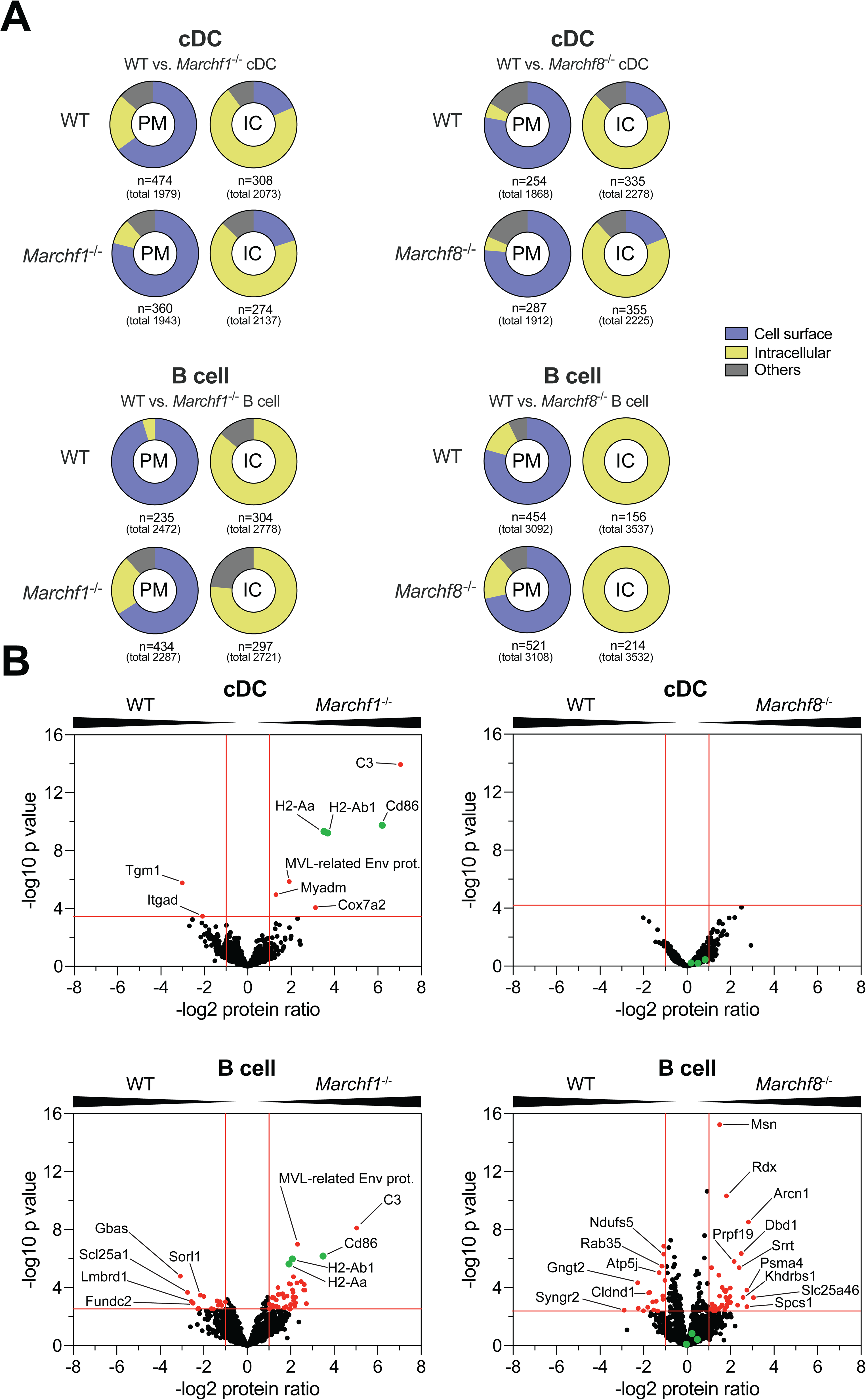
Proteomic analysis of differentially expressed proteins in the plasma membrane fraction between WT and *Marchf1^−/−^* or *Marchf8^−/−^* cDC and B cells. Proteomic analysis of plasma membrane (PM)-enriched microsome fractions of splenic cDC and B cells purified from WT, *Marchf1*^−/−^ or *Marchf8*^−/−^ mice. PM fractions were purified from post-nuclear supernatants of mAb surface stained cDC and B cells via magnetic immunoaffinity and analysed by semi-quantitative mass spectrometry from three biological replicates (in total 3x 8 samples; WT vs. *Marchf1*^−/−^ and WT vs. *Marchf8*^−/−^ cDC + WT vs. *Marchf1*^−/−^ and WT vs. *Marchf8*^−/−^ B cells). The remaining compartments (mitochondria, endosomes, etc.) from the post-nuclear supernatant of homogenized cells following PM fraction retrieval was termed intracellular compartment (IC). (A) Enrichment analysis (performed using the function enricher included in the Bioconductor clusterProfiler package [42]) of detected proteins via MS from PM or IC fractions from cDC and B cells of WT versus *Marchf1*^−/−^ and WT versus *Marchf8*^−/−^ mice. Annotated GO-IDs for detected proteins were grouped into categories of ‘Cell surface’, ‘Intracellular Compartment (IC)’ and ‘others’ based on experimentally verified Gene Ontology (GO) annotations. ‘Cell surface’ category included the GO terms ‘plasma membrane’, ‘external side of plasma membrane’ and ‘cell surface’ among others, while the categories ‘Intracellular Compartment (IC)’ and ‘others’ included GO terms such as ‘mitochondrial membrane’ and ‘endoplasmic reticulum’ as well as ‘myelin sheath’, respectively. For a detailed list of all annotated GO terms of all fractions please see Supplementary Figure 4. (B) Detection of differentially expressed proteins in the PM fraction of cDC and B cells of WT versus *Marchf1*^−/−^ and WT versus *Marchf8*^−/−^ mice. Equivalent amounts of PM fractions (based on cell count) of three biological replicates were analyzed by mass spectrometry and semi-quantitative proteomics in three technical replicates. Proteins detected in both WT and *Marchf1*^−/−^ or *Marchf8*^−/−^ cDC/B cells were displayed in volcano plots (1020 proteins for WT vs. *Marchf1*^−/−^ cDC, 922 proteins for WT vs. *Marchf8*^−/−^ cDC, 1275 proteins for WT vs. *Marchf1*^−/−^ B cells and 1819 proteins for WT vs. *Marchf8*^−/−^ B cells) with differentially expressed proteins [red dots] identified based on two-fold ratio (log2 protein ratio >1 or <1) and significance (5% FDR) across three biological replicates, each measured in technical triplicates. The known MARCH1 substrates, MHC II (H2-Aa and H2-Ab1) and CD86 in B cells and cDC, are highlighted in green in each volcano plot.

Comparison of the PM proteomes of WT and *Marchf1*^−/−^ cDC showed that, as expected, most proteins were present at similar levels in the two preparations (1020 proteins in total, **Supplementary Table 3)**. Nine proteins were differentially expressed between WT and *Marchf1*^−/−^ cDC PM [log2 protein ratio >1 or <1 and -log10 adjusted *p* value >3.47 (5% FDR)] (**Figure 5B** and **Supplementary Table 5**). These included MHC IIα and β chains (H2-Aa and H2-Ab1), as well as CD86, confirming the validity of our approach to detect MARCH1 substrates. Surprisingly, the protein that appeared most significantly overexpressed in the PM of *Marchf1*^−/−^ cDC was complement component 3 (C3) (**Figure 5B, Supplementary Table 5**). The remaining three proteins appearing over-expressed in the *Marchf1*^−/−^ cDC PM fraction are not known to be immunoreceptors expressed at the PM: Cox7a2 is a mitochondrial protein, Myadm a component of the cytoskeleton and MLV-related proviral Env polyprotein, a protein endogenously encoded by a retrovirus integrated in the genome of commonly used mouse strains [58]. As our main goal was to identify immunoregulatory MARCH1 substrates, we did not investigate further whether these were true or artifactual “hits” of the proteomic analysis. Comparison of the PM fractions of WT and *Marchf8*^−/−^ cDC did not reveal any differentially expressed proteins (**Figure 5B,** 922 proteins in total, **Supplementary Table 3**), supporting the previous results indicating that MARCH8 is not expressed/active in cDC.

*Marchf1*^−/−^ and *Marchf8*^−/−^ B cells exhibited 45 and 40 enriched and 15 and 17 reduced proteins, respectively, in their PM fractions [log2 protein ratio >1 or <1. and -log10 adjusted *p* value >2.5 and >2.36 for *Marchf1*^−/−^ and *Marchf8*^−/−^, respectively (both 5% FDR)] (**Figure 5B, Supplementary Table 6** and **Supplementary Table 7**, 1275 and 1819 proteins in total, **Supplementary Table 3**). MHC IIα and β chains (H2-Aa and H2-Ab1), as well as CD86 and C3 were the most significantly enriched proteins in the PM fraction of *Marchf1*^−/−^ B cells (**Figure 5B** and **Supplementary Table 6**), but neither of the four were enriched in *Marchf8*^−/−^ B cells (**Figure 5B** and **Supplementary Table 7)**. Only 14 of the 60 proteins differentially expressed in the PM fraction of *Marchf1*^−/−^ B cells, and 10 of the 57 proteins differentially expressed in the PM fraction of *Marchf8*^−/−^ B cells, were immunoreceptors and/or proteins known to be expressed at the plasma membrane (**Supplementary Table 6** and **Supplementary Table 7**). They included aminopeptidase N (CD13, gene *Anpep*), antigen-presenting glycoprotein CD1d, T cell differentiation antigen CD6 and the immunoglobulin epsilon Fc receptor CD23 (gene *Fcer2*). However, analysis by flow cytometry did not confirm differential expression in either *Marchf1*^−/−^ or *Marchf8*^−/−^ B cells (**Supplementary Figure 5).** The most likely explanation for detection of these “false positives” is that they were caused by subtle differences in the purity of the B cell preparations or their subcellular fractions. In conclusion, MHC class II and CD86 were the only membrane proteins that we could unequivocally confirm as MARCH1 substrates in B cells, and while we cannot discard the possibility that some of the “hits” found in the proteomic screen of *Marchf8*^−/−^ B cells are indeed MARCH8 substrates, it is more likely that MARCH8 is not active in B cells, just as it is not in DC.

### MARCH8, not MARCH1, is active in non-hematopoietic cells

The only cell type in which MARCH8 activity has been demonstrated is thymic epithelial cells (TEC), where it regulates MHC II surface expression but not CD86 [9][10]. Analysis of CD40, CD44, CD95 and CD98 expression in WT and *Marchf8*^−/−^ medullar and cortical TEC showed that neither of these receptors, which have been shown to be ubiquitinated in cell lines overexpressing MARCH8, are physiological substrates (**Figure 6A**).

**Figure 6.**
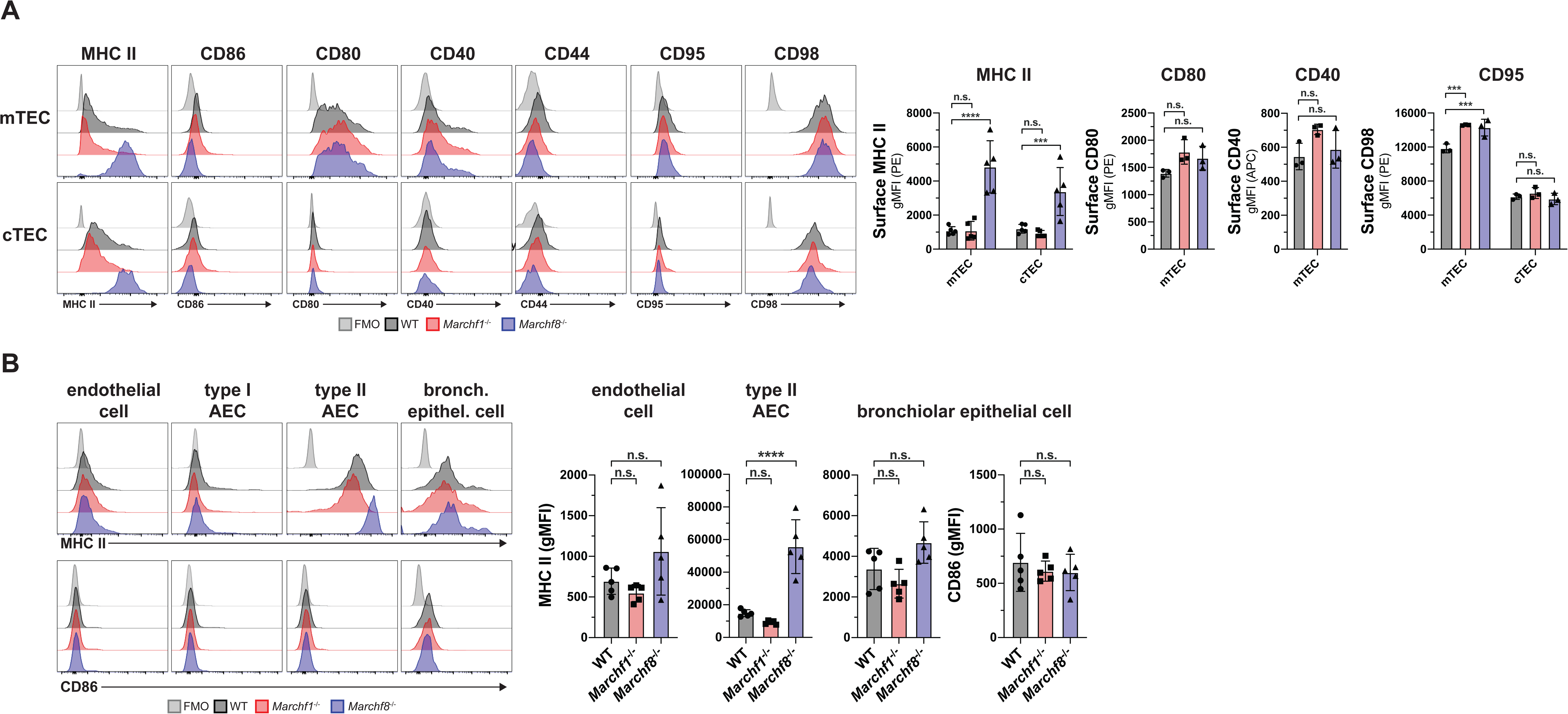
Ubiquitination of MHC II, CD86 and putative substrates by MARCH1 and MARCH8 in non-haemopoietic antigen presenting cells. (A) Surface expression of MHC II, CD86, CD80, CD40, CD44, CD95 and CD98 in medullary and cortical thymic epithelial cells (mTEC and cTEC) purified from WT, *Marchf1*^−/−^ and *Marchf8^−/−^* mice. (B) Surface expression of MHC II and CD86 in endothelial cells, type I and type II alveolar epithelial cells (AEC) as well as bronchiolar epithelial cells, purified from the lung of WT, *Marchf1*^−/−^ and *Marchf8^−/−^* mice. In all cases a fluorescence-minus-one (FMO) control was included, for which cells were incubated with the corresponding multi-colour staining panel, excluding the fluorescently labelled antibody species of interest. Bars represent mean ± SD with each symbol representing an individual mouse (n=5). Statistical analysis was performed using one-way ANOVA followed by Sidak’s multiple comparisons test. **** p < 0.0001, *** p < 0.0002, ** p < 0.002, * p < 0.03, n.s. not significant.

Although TEC constitutively present antigens via MHC II, they are not hematopoietic cells, but of endodermal origin [59]. Therefore, we asked the question whether other cells ontogenically related to TEC also use MARCH8 to regulate surface MHC II expression. Epithelial cells in the respiratory tract are known to express MHC II, with the highest level found on type II alveolar epithelial cells (AEC) [60][61][62]. We found that MARCH8- deficient type II AEC showed enriched MHC II surface expression (**Figure 6B**), but MHC II levels in mutant endothelial cells, type I AEC and bronchial epithelial cells was not altered (**Figure 6B**). Neither cell type displayed increased CD86 expression in the absence of MARCH8, and lack of MARCH1 did not affect MHC II nor CD86 expression in any of the cell types analyzed (**Figure 6B**). In conclusion, not all epithelial cells regulate MHC II expression via ubiquitination, but those that do employ MARCH8.

## DISCUSSION

Determining which cells utilize MARCH1 and MARCH8 has been hampered by their low level of expression, but analysis of MHC II and CD86 as surrogate markers of activity has allowed us to establish the role of MARCH1 as a master regulator of MHC II and CD86 expression in all hematopoietic cells. MARCH8 plays an equivalent role in the two major types of TEC and in type II AEC, where it ubiquitinates MHC II. We did not observe high CD86 expression in any *Marchf8*^−/−^ cell, but this could be because these cells do not ubiquitinate CD86 or because they do not express it. There are at least two precedents for ontogeny-specific differences in the use of components of MHC II antigen presentation machinery. Expression of CIITA, which directs transcription of the genes for MHC II and for several accessory molecules involved in antigen presentation, is driven by distinct promoters in hematopoietic and non-hematopoietic cells [63]. Proteolysis of the chaperone invariant chain, a critical step in the MHC II antigen presentation pathway, is carried out by cathepsin S in hematopoietic cells and by and cathepsin L in non-hematopoietic cells [64]. It is unclear why this dichotomy exists, which is probably caused by the establishment of cell lineage-specific gene programs during embryonic development.

While our finding that MARCH1 is operative in professional APC confirmed previous observations, we were surprised to observe high MHC II and CD86 expression in non-professional APC lacking MARCH1. This was not caused by ectopic induction or overexpression of either molecule because MARCH1-deficient T cells maintained WT levels of MHC II (negative) and CD86 (low) expression. As MARCH1 ubiquitinates substrates that have already trafficked through the cell surface, this finding implies that atypical APC express and deposit on their plasma membrane larger amounts of MHC II and CD86 than is usually appreciated, but their steady-state levels are kept low by virtue of MARCH1 ubiquitination and accelerated turn-over. Eosinophils are associated with inflammatory responses during allergy or parasitic infections, while neutrophils are recruited in abundant numbers to sites of tissue damage or infection. The role of MHC II antigen presentation by either cell type is controversial. While there is evidence for both purified eosinophils and neutrophils that demonstrates their capacity to present antigen via MHC II [46], it is difficult to exclude the possibility of DC contamination in these assays. *In vivo* evidence of their antigen presentation capacity is scarce but there are reported examples where both eosinophils [65] and neutrophils [66][67][68] contribute to enhancing antigen-specific CD4^+^ T cell responses. The realization that these cells regulate MHC II and CD86 via ubiquitination utilizing the same mechanism as professional APC lends weight to the notion that they perform antigen presentation *in vivo*.

One of the functions attributed to MARCH8 in humans is to ubiquitinate viral proteins deposited on the plasma membrane of infected cells and that will be incorporated in the envelop of the virion upon budding [69][70][71]. The reduction of viral protein expression that ensues inhibits spread of the infection, protecting the host. This activity has not been described in mice, but our results suggest that if it occurs in this species, it is unlikely to be operative in hematopoietic cells, where perhaps other members of the MARCH family replace the function of MARCH8.

While several substrates have been identified for MARCH1 and MARCH8 based on studies using overexpression and/or cell lines, our flow cytometry analysis rules out CD44, CD71, CD95 and CD98 as bona fide MARCH1 or MARCH8 substrates in all primary cells examined. This highlights that caution needs to be taken when interpreting studies that rely on E3 Ub ligase overexpression. Our unbiased proteomic profiling of B cells and DC unequivocally confirmed the role of MARCH1 in MHC II and CD86 ubiquitination in both cell types, but did not reveal any other MARCH1 substrate that we could validate by flow cytometry with the exception of complement C3. Further investigations will be required to determine if enriched levels of surface C3 in these cells is a direct or indirect effect of MACRH1 deficiency, as we have also shown that high MHC II expression in *Marchf1*^−/−^ cells indirectly induces higher or lower expression of other surface receptors that are not direct MARCH1 substrates [72]. However, the magnitude of these changes is below the level of resolution afforded by high-throughput, unbiased proteomic analysis of subcellular fractions. We did not observe changes in expression of any protein on the plasma membrane of *Marchf8*^−/−^ DC. The “hits” detected *Marchf8*^−/−^ B cell membrane could be attributed to contamination with other subcellular compartments because they were not classified as plasma membrane proteins and/or could not be validated as differentially expressed by flow cytometry. The proteomic analysis thus confirmed that neither B cells nor DC express functional MARCH8.

In summary, MHC II is the only membrane protein unequivocally regulated by MARCH1 and MARCH8 in primary mouse cells, with each ligase playing its role in haemopoietic and non-haemopoietic cells, respectively. CD86 is also a MARCH1 substrate in hematopoietic cells.

These results help to predict the potential effects of genetic or pharmacological manipulation of MARCH1 or MARCH8 activities as a treatment for immunological disorders.

## ACKNOWLEDGEMENTS

We thank the Antibody Services Facility and Genomics Hub (Walter and Eliza Hall Institute) and the Melbourne Cytometry Platform (The University of Melbourne) for expert assistance. JAV: NHMRC Fellowship 1058193 and 1154502, NHMRC Program 1016629 and 1113293, ARC DP160103134 and DP110101383, and Human Frontiers Science Program Grant 0064/2011. PS: Australian Research Training Programme Scholarship provided by the Australian Commonwealth Government and the University of Melbourne.

## SUPPLEMENTARY FIGURE LEGENDS

**Supplementary Figure 1.**
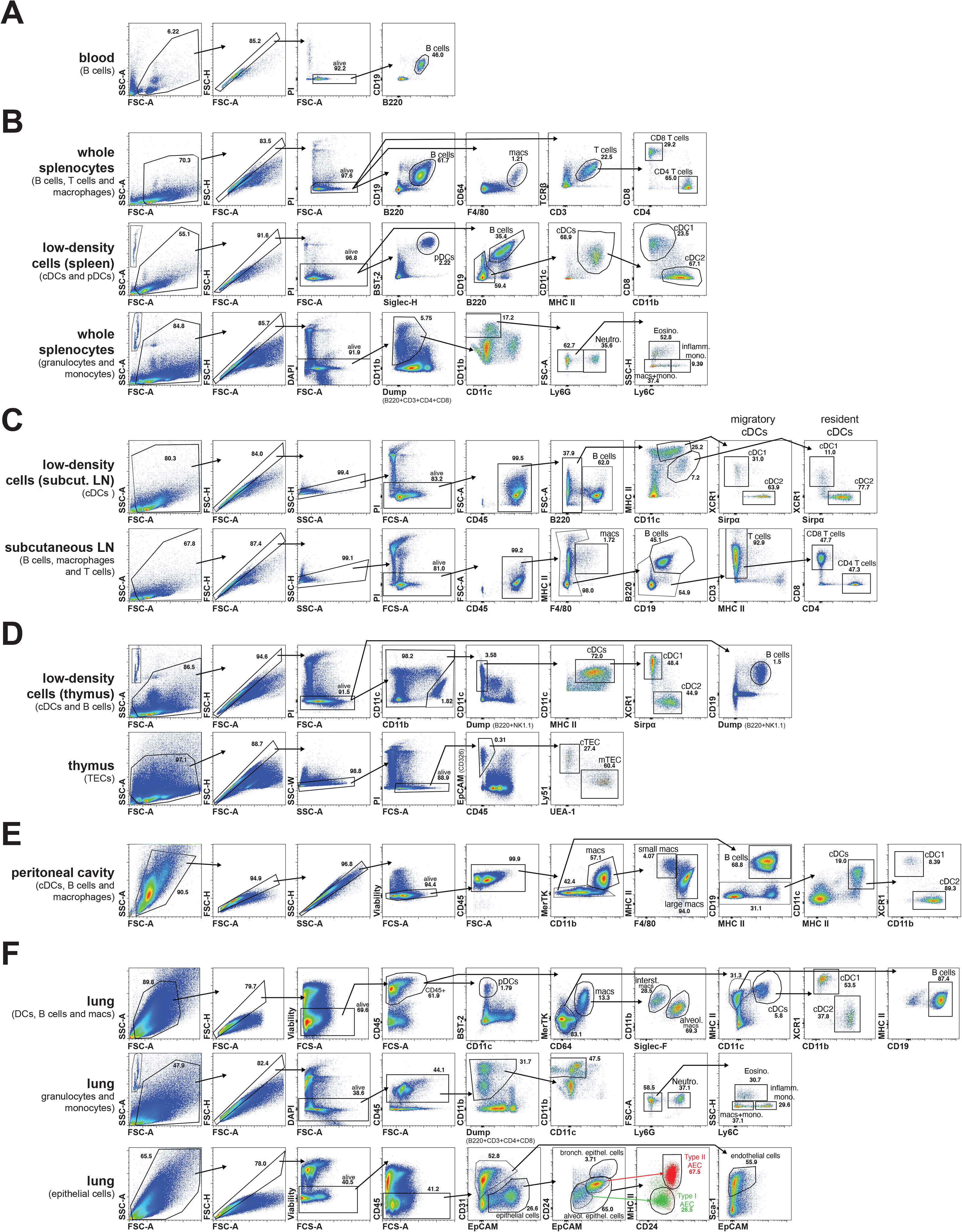
Representative flow cytometry gating strategies for the identification of cell populations of interest in blood, spleen, subcutaneous lymph nodes (LN), thymus, peritoneal cavity and lung. In all cases cell doublets and dead cells were identified and excluded based on forward and side scatter (FSC and SSC) as well as staining with propidium iodide (PI), diamidino phenylindole (DAPI) or Fixable Viability Dye eFluor™780 (Viability). (A) Blood B cells were identified as CD19^+^ B220^+^. (B) Splenic B cells, macrophages and T cells were identified from whole splenocyte suspensions as CD19^+^ B220^+^, F4/80^+^ CD64^+^ and TCRβ^+^ CD3^+^ respectively with further discrimination of CD4^+^ and CD8^+^ T cells. Splenic DC were identified from low-density splenocyte suspensions, with pDC identified as Siglec-H^+^ BST-2^+^ and cDC as B220^−^ CD19^−^ CD11c^+^ MHC II^+^, with further discrimination of cDC1 as CD11b^−^ CD8^+^ and cDC2 as CD11b^+^ CD8^−^. Splenic granulocytes and monocytes were identified from whole splenocyte suspensions as B220^−^ CD3^−^ CD4^−^ CD8^−^ CD11c^low-mid^ CD11b^high^ with neutrophils identified as Ly6G^+^, eosinophils as Ly6G^−^ SSC-H^high^ Ly6C^low-mid^, patrolling monocytes as Ly6G^−^ Ly6C^low- mid^ SSC-H^low^ and inflammatory monocytes as Ly6G^−^ Ly6C^high^ SSC-H^low^ (as described in Liyanage *et al*. [73]). (C) cDC from subcutaneous LN were identified from low-density cell suspensions, with resident cDC identified as CD11c^high^ MHC II^mid^ and migratory cDC identified as CD11c^mid^ MHC II^high^ and further discrimination of cDC1 as Sirpα^−^ XCR1+ and cDC2 as Sirpα^+^ XCR1^−^. B cells, macrophages and T cells from subcutaneous LN were identified from whole cell suspensions as CD19^+^ B220^+^, F4/80^+^ MHC II^+^ and CD3^+^ respectively with further discrimination of CD4^+^ and CD8^+^ T cells. (D) Thymic cDC were identified from low-density cell suspensions as B220^−^ NK1.1^−^ CD11c^+^ MHC II^+^, with further discrimination of cDC1 as Sirpα^−^ XCR1^+^ and cDC2 as Sirpα+ XCR1- (as described in Ardouin *et al*. [74]). Thymic epithelial cells (TECs) were identified from whole thymocyte suspensions as CD45^−^ EpCAM^+^, with further discrimination of cortical TECs (cTECs) as UEA-1^−^ Ly51^+^ and medullary TECs (mTECs) as UEA-1^+^ Ly51^−^ (as described in Liu *et al*. [9]). (D) Peritoneal macrophages were identified as CD11b^+^ MerTK^+^, with further discrimination of small peritoneal macrophages as F4/80^low^ MHC II^high^ and large peritoneal macrophages as F4/80^high^ MHC II^mid-high^ (as described in Bain *et al*. [75]). Peritoneal B cells were identified as MerTK^−^ MHC II^+^ CD19^+^ and cDC as MerTK^−^ CD19^−^ CD11c^+^ MHC II^+^ with further discrimination of cDC1 as CD11b^−^ XCR1^+^ and cDC2 as CD11b^+^ XCR1^−^. (E) Haemopoietic cells in the lung were identified as CD45^+^ with pDC as CD11c^low-mid^ BST-2^+^ and macrophages as CD64^+^ MerTK^+^, with further discrimination of interstitial macrophages as Siglec-F^low^ CD11b^high^ and alveolar macrophages as Siglec-F^high^ CD11b^mid^ (as described in Svedberg *et al*. [76]). Lung B cells were identified as CD64^−^ MerTK^−^ CD11c^−^ MHC II^+^ CD19^+^ and cDC as CD64^−^ MerTK^−^ CD11c^+^ MHC II^+^ with further discrimination of cDC1 as CD11b^−^ XCR1^+^ and cDC2 as CD11b^+^ XCR1^−^. Lung granulocytes and monocytes were identified as B220^−^ CD3^−^ CD4^−^ CD8^−^ CD11c^low-mid^ CD11b^high^ with neutrophils identified as Ly6G^+^, eosinophils as Ly6G^−^ SSC-H^high^ Ly6C^low-mid^, patrolling monocytes as Ly6G^−^ Ly6C^low-mid^ SSC-H^low^ and inflammatory monocytes as Ly6G^−^ Ly6C^high^ SSC-H^low^ (as described in Liyanage *et al*. [73]). Non-haemopoietic cells in the lung were identified as CD45^−^ with endothelial cells identified as EpCAM^low-mid^ CD31^+^ Sca-1^+^ and epithelial cells as EpCAM^mid-high^ CD31^−^. Further discrimination of epithelial cells was carried out based of CD24, EpCAM and MHC II expression (as described in Nakano *et al*. [61] and Hasegawa *et al*. [62]) with bronchiolar epithelial cells identified as EpCAM^high^ CD24^high^, type II alveolar epithelial cells (AEC) identified as EpCAM^high^ CD24^mid^ MHC II^high^ (red) and type I AEC identified as EpCAM^mid^ CD24^low^ MHC II^low^ (green). A comparison of the representative flow cytometry gating strategies for the identification of all cell populations of interest between WT, *Marchf1*^−/−^ and *Marchf8*^−/−^ mice is shown in Supplementary Figure 2.

**Supplementary Figure 2.**
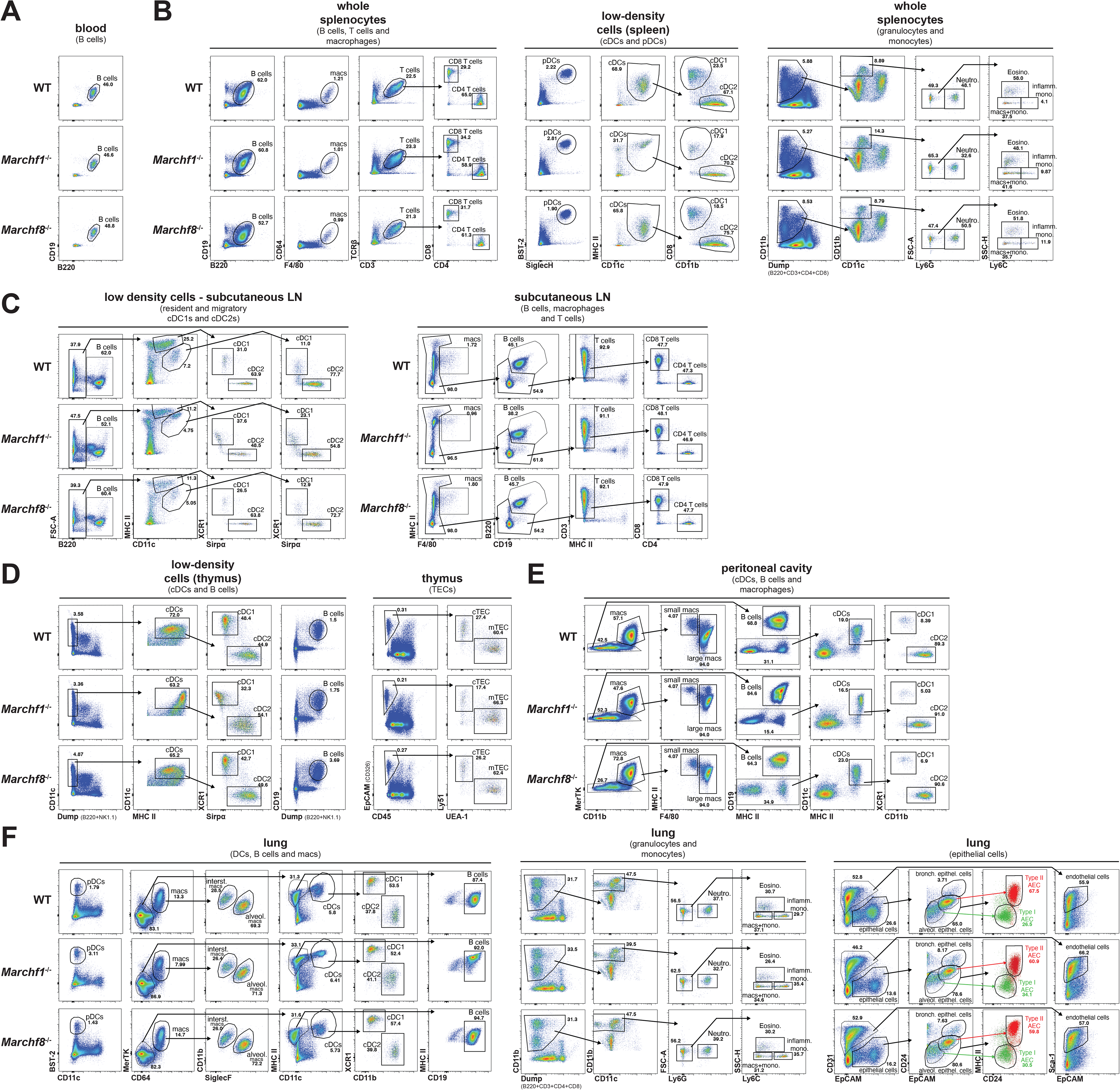
Comparison of representative flow cytometry gating strategies for the identification of various cell populations in (A) blood, (B) spleen, (C) subcutaneous lymph node, (D) thymus, (E) peritoneal cavity and (F) lung from WT, *Marchf1*^−/−^ and *Marchf8*^−/−^ mice. A detailed description of the gating strategies for each individual cell population of interest is presented in Supplementary Figure 1.

**Supplementary Figure 3.**
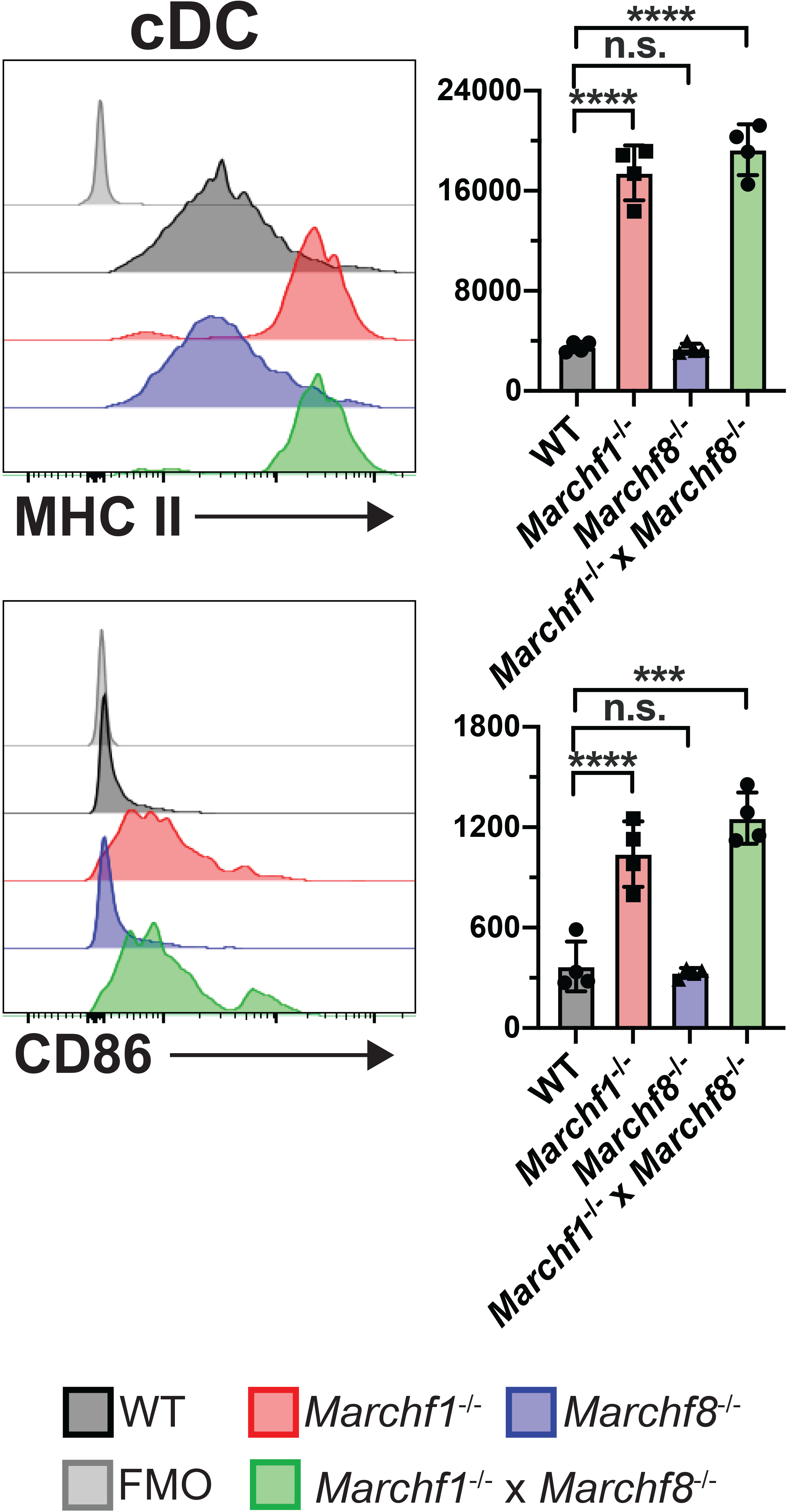
Surface expression of MHC II and CD86 in peritoneal cDC from WT, *Marchf1*^−/−^ and *Marchf8*^−/−^ mice or from mice deficient in both MARCH1 and MARCH8 (*Marchf1*^−/−^ x *Marchf8*^−/−^). A fluorescence-minus-one (FMO) control was included, for which cells were incubated with the corresponding multi-colour staining panel, excluding the fluorescently labelled antibody species of interest (i.e. anti-CD86 or anti-MHC II mAb). Bars represent mean ± SD with each symbol representing an individual mouse (n=4). Statistical analysis was performed using one-way ANOVA followed by Sidak’s multiple comparisons test. **** p < 0.0001, *** p < 0.0002, ** p < 0.002, * p < 0.03, n.s. not significant.

**Supplementary Figure 4.**
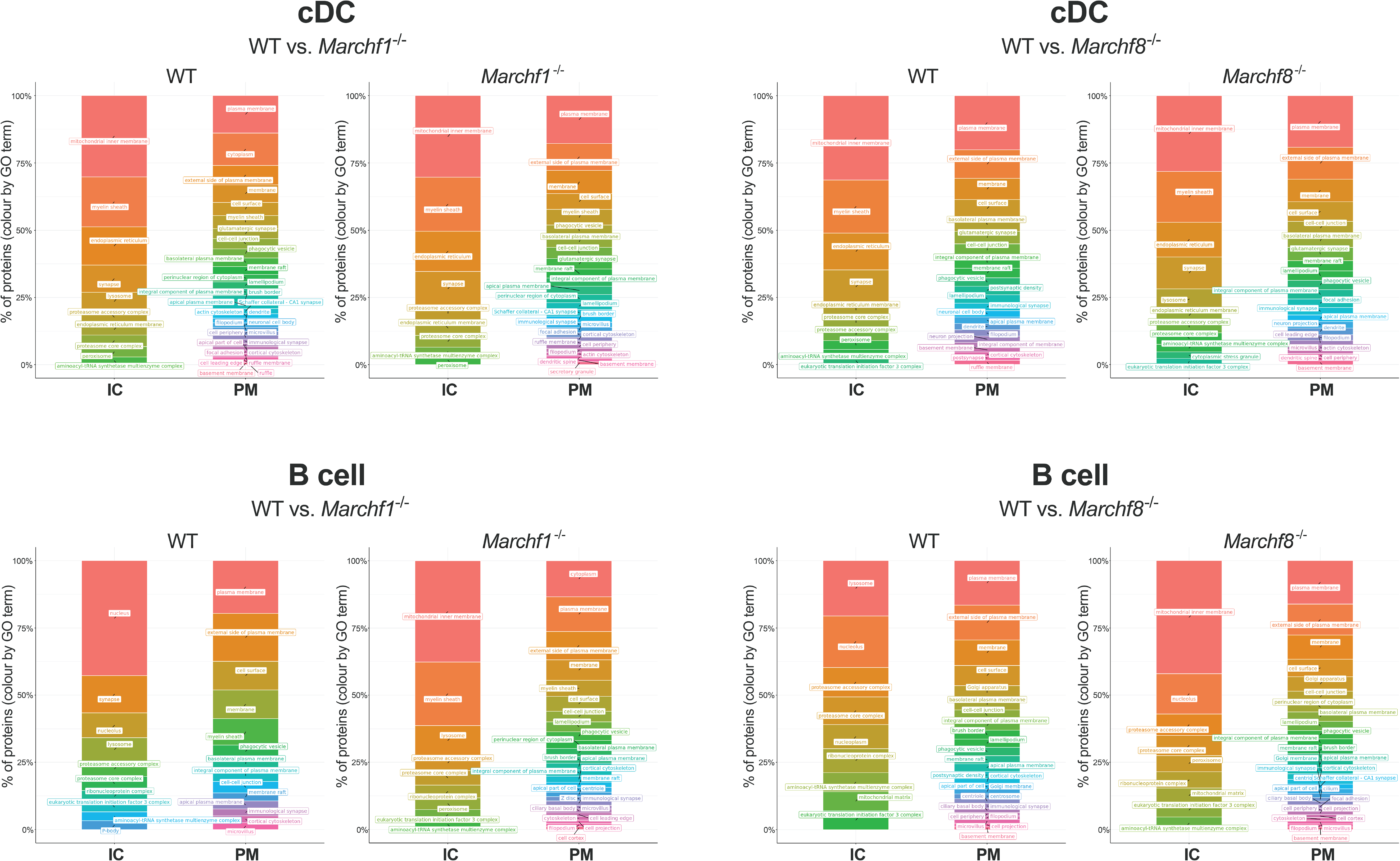
Gene ontology (GO) enrichment analysis of proteins detected in plasma membrane (PM)- enriched and intracellular compartment (IC)-enriched microsome fractions of splenic cDC and B cells purified from WT, *Marchf1*^−/−^ or *Marchf8*^−/−^ mice. PM fractions were purified from post- nuclear supernatants of mAb surface stained cDC and B cells via magnetic immunoaffinity. IC (intracellular compartments) was retrieved from the post-nuclear supernatant of homogenized cells following PM fraction extraction. IC and PM fraction were analysed by semi-quantitative mass spectrometry and GO term enrichment analysis was performed using the Bioconductor clusterProfiler package [42] with GO-IDs grouped based on experimentally verified Gene Ontology (GO) annotations.

**Supplementary Figure 5.**
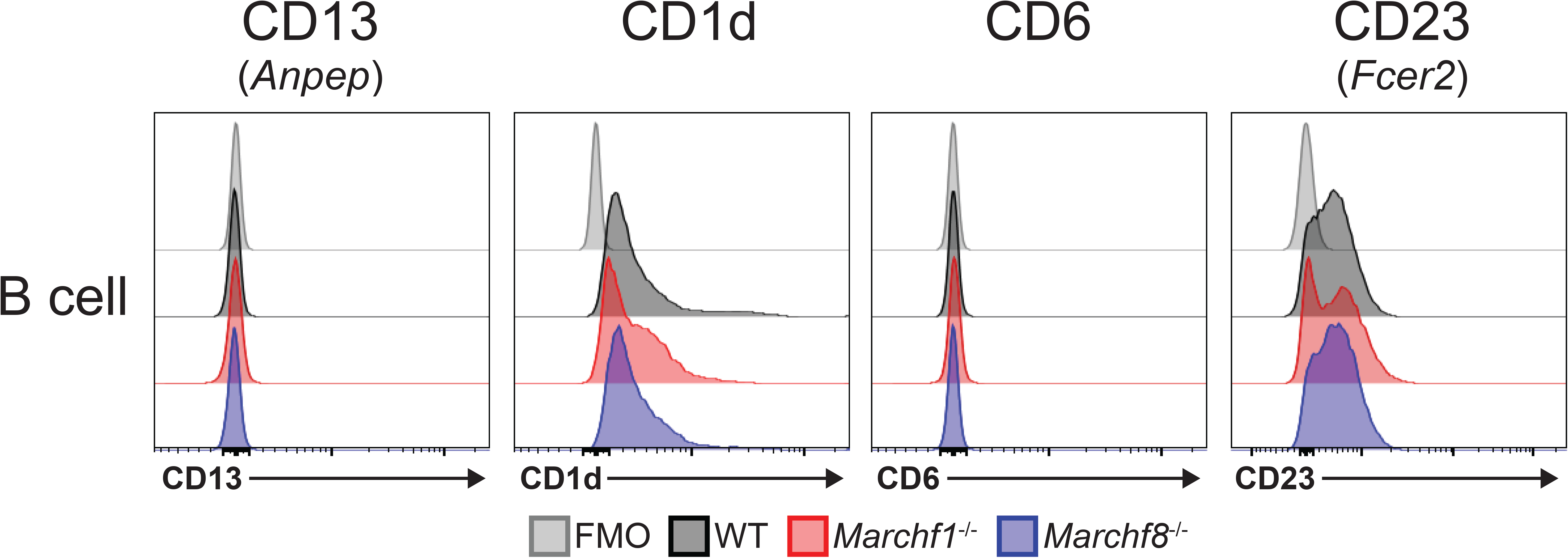
Surface expression of CD13, CD1d, CD6 and CD23 in B cells from WT, *Marchf1*^−/−^ and *Marchf8*^−/−^ mice. A fluorescence-minus-one (FMO) control was included, for which cells were incubated with the corresponding multi-colour staining panel, excluding the fluorescently labelled antibody species of interest.

**Supplementary Table 5:** Significantly up/down-regulated proteins in the PM fraction of *Marchf1*^−/−^ cDCs.

| **Gene** | **Protein Names** | **Log2 Fold Change** | **-Log10 p value** | **Localisation (based on GO-ID of enrichment analysis)** | **Localisation (based on UniProt)** |
| --- | --- | --- | --- | --- | --- |
| ***C3*** | Complement C3 | 7.04 | 13.95 | Extracellular, Cell surface | Extracellular region or secreted |
| ***Cd86*** | T-lymphocyte activation antigen CD86 | 6.20 | 9.76 | Plasma membrane, Intracellular membrane-bounded organelle | Cell membrane, Single-pass type I membrane protein |
| ***H2-Aa*** | H-2 class II histocompatibility antigen, A-B alpha chain | 3.51 | 9.33 | Plasma membrane, Early endosome | Membrane, Single-pass type I membrane protein |
| ***H2-Ab1*** | H-2 class II histocompatibility antigen, A beta chain | 3.68 | 9.22 | Plasma membrane, Lysosome | Membrane, Single-pass type I membrane protein |
| **-** | MLV-related proviral Env polyprotein | 1.91 | 5.85 | N/A | Cell membrane, Virion membrane |
| ***Myadm*** | Myeloid-associated differentiation marker | 1.29 | 4.95 | N/A | Cortical actin cytoskeleton, Membrane, Multi-pass membrane protein |
| ***Cox7a2*** | Cytochrome c oxidase subunit 7A | 3.12 | 4.06 | Mitochondrial inner membrane | Mitochondrion inner membrane |
| ***Tgm1*** | Protein-glutamine gamma-glutamyltransferase K | -3.02 | 5.77 | Adherens junction | Membrane, Lipid-anchor |
| ***Itgad*** | Integrin alpha-D | -2.08 | 3.47 | N/A | Membrane, Single-pass type I membrane protein |

**Supplementary Table 6:** Significantly up/down-regulated proteins in the PM fraction of *Marchf1*^−/−^ B cells.

| Gene | Protein Names | Log2 Fold Change | -Log10 p value | Localisation<br>(based on GO-ID of enrichment analysis) | Localisation<br>(based on UniProt) |
| --- | --- | --- | --- | --- | --- |
| **C3** | Complement C3 | 5.04 | 8.12 | Extracellular, Cell surface | Extracellular region or secreted |
| **-** | MLV-related proviral Env polyprotein | 2.32 | 6.99 | N/A | Cell membrane, Virion membrane |
| **Cd86** | T-lymphocyte activation antigen CD86 | 3.49 | 6.18 | Plasma membrane & intracellular membrane-bounded organelle | Cell membrane, Single-pass type I membrane protein |
| **H2-Ab1** | H-2 class II histocompatibility antigen, A beta chain | 2.07 | 5.98 | Plasma membrane, Lysosome | Membrane, Single-pass type I membrane protein |
| **H2-Aa** | H-2 class II histocompatibility antigen, A-B alpha chain | 1.92 | 5.64 | Plasma membrane, Early endosome | Membrane, Single-pass type I membrane protein |
| **Stk24; Stk25; Stk26** | Serine/threonine-protein kinase 24/25/26 | 2.13 | 4.76 | N/A | Nucleus |
| **Anpep** | Aminopeptidase N | 2.48 | 4.42 | Plasma membrane | Cell membrane, Single-pass type II membrane protein |
| **Ahsg** | Alpha-2-HS-glycoprotein | 2.29 | 4.32 | N/A | Secreted |
| **Lrch1** | Leucine-rich repeat and calponin homology domain-containing protein 1 | 1.94 | 4.27 | N/A | Cytoplasm |
| **Ttc7a** | Tetratricopeptide repeat protein 7A | 1.99 | 4.25 | N/A | Cell membrane, Cytoplasm |
| ***Rpl34*** | 60S ribosomal protein L34 | 2.63 | 4.22 | Mitochondrion | Endoplasmic reticulum,<br>Cytosol |
| ***Coro2a*** | Coronin-2A | 1.42 | 4.00 | Brush border | Brush border,<br>Transcription repressor |
| ***Sh3kbp1*** | SH3 domain-containing kinase-binding protein 1 | 2.08 | 3.84 | Cell-cell junction,<br>Endocytic vesicle | Cytoskeleton,<br>Cytoplasm |
| ***Ifi30*** | Gamma-interferon-inducible lysosomal thiol reductase | 2.60 | 3.83 | Lysosome | Lysosome |
| ***Itgax*** | Integrin alpha-X | 2.65 | 3.83 | Plasma membrane | Membrane, Single-pass<br>type I membrane protein |
| ***Hbb-b1;Hbb-b2*** | Hemoglobin subunit beta-1/2 | 2.11 | 3.68 | Myelin sheath,<br>Haemoglobin complex | Cytosol |
| ***Stk10*** | Serine/threonine-protein kinase 10 | 1.63 | 3.58 | N/A | Cell membrane,<br>Peripheral membrane |
| ***Frmd8*** | FERM domain-containing protein 8 | 1.56 | 3.55 | N/A | Cytosol, Cell membrane |
| ***Taok3*** | Serine/threonine-protein kinase TAO3 | 1.43 | 3.53 | N/A | Cytoplasm |
| ***Git2*** | ARF GTPase-activating protein GIT2 | 1.83 | 3.42 | Calyx of Held | Nucleoplasm |
| ***Actr2*** | Actin-related protein 2 | 1.18 | 3.36 | Cell cortex, Actin cap | Cytoskeleton, Nucleus |
| ***Ptprj*** | Receptor-type tyrosine-protein phosphatase eta | 2.10 | 3.34 | Immunol. synapse,<br>Plasma membrane,<br>Ruffle membrane | Cell membrane, Single-pass<br>type I membrane protein |
| ***Ahrr*** | Aryl hydrocarbon receptor repressor | 2.23 | 3.34 | Nucleus | Nucleus |
| ***Fam126a*** | Hyccin | 1.69 | 3.31 | Neuron projection | Cytosol, Plasma<br>membrane |
| ***Stxbp3*** | Syntaxin-binding protein 3 | 1.16 | 3.29 | Plasma membrane,<br>Apical plasma<br>Membrane, Cytosol | Cell membrane, Cytosol |
| ***Gbp5*** | Guanylate-binding protein 5 | 1.24 | 3.28 | Cytoplasmic vesicle | Golgi apparatus<br>membrane, Cytoplasm |
| ***Stxbp2*** | Syntaxin-binding protein 2 | 1.34 | 3.15 | Apical plasma membrane, Phagocytic vesicle, Zymogen granule membrane | Cytosol, azurophil granule, apical plasma membrane |
| ***Cct8*** | T-complex protein 1 subunit theta | 2.13 | 3.12 | Cell body, Zona pellucida receptor complex, Chaperonin-containing T-complex | Cytoskeleton, Cytoplasm |
| ***Ap1m1*** | AP-1 complex subunit mu-1 | 2.23 | 2.96 | N/A | Golgi apparatus, Peripheral membrane protein |
| ***Pacsin2*** | Protein kinase C and casein kinase substrate in neurons protein 2 | 2.72 | 2.90 | Cytoplasm, Cell-cell junction, Cytosol, Trans-Golgi network, Extrinsic component of membrane | Ruffle membrane. Peripheral membrane, Cell membrane, Early endosome, Cytoskeleton |
| ***Iqgap1*** | Ras GTPase-activating-like protein IQGAP1 | 1.00 | 2.89 | Nucleus, Cytoplasm, Cell-cell junction, Lateral plasma membrane, Neuron projection, Cell leading edge, Ribonucleoprotein complex | Nucleus, Plasma membrane, Cytoplasm |
| ***Agfg1*** | Arf-GAP domain and FG repeat-containing protein 1 | 2.18 | 2.85 | Cytoplasmic vesicle, Neuronal cell body, Cell projection | Nucleus, Cytoplasmic vesicle |
| ***Hba*** | Hemoglobin subunit alpha | 2.00 | 2.83 | Myelin sheath | Cytosol, Extracellular region or secreted, Myelin sheath |
| ***Tubgcp3*** | Gamma-tubulin complex component 3 | 1.16 | 2.81 | N/A | Centrosome |
| ***Csk*** | Tyrosine-protein kinase CSK | 1.16 | 2.77 | Cell-cell junction | Plasma membrane, Cytoplasm |
| ***Ap1b1*** | AP-1 complex subunit beta-1 | 2.19 | 2.74 | N/A | Golgi apparatus,<br>Peripheral membrane<br>protein |
| ***Actr3*** | Actin-related protein 3 | 1.14 | 2.73 | Lamellipodium, Cell-cell<br>junction, Brush border | Cytoskeleton, Nucleus,<br>Cell projection |
| ***Ablim1*** | Actin-binding LIM protein 1 | 1.22 | 2.73 | Actin cytoskeleton,<br>Postsynaptic density | Cytoskeleton,<br>Cytoplasm |
| ***Ap2s1*** | AP-2 complex subunit sigma | 1.35 | 2.71 | AP-2 adaptor complex | Cell membrane,<br>Peripheral membrane<br>protein |
| ***Eps15*** | Epidermal growth factor receptor substrate<br>15 | 1.79 | 2.68 | Plasma membrane,<br>Clathrin-coated<br>pit/vesicle, Ciliary<br>membrane, AP-2<br>adaptor complex | Cell membrane,<br>Peripheral membrane<br>protein, Cytoplasm,<br>Clathrin-coated pit |
| ***Ccm2*** | Cerebral cavernous malformations protein<br>2 homolog | 1.43 | 2.66 | Protein-containing<br>complex | Cytoplasm |
| ***Fam65b*** | Protein FAM65B | 1.70 | 2.60 | Stereocilium | Cytoskeleton,<br>Stereocilium membrane,<br>Apical cell membrane,<br>Cytoplasm |
| ***Arpc2*** | Actin-related protein 2/3 complex subunit 2 | 1.02 | 2.53 | Focal adhesion, Plasma<br>membrane, Synapse,<br>Endosome, Cell leading<br>edge, | Nucleus, Cytoskeleton,<br>Cell projection |
| ***Anxa6*** | Annexin A6 | 1.99 | 2.52 | Perinuclear region of<br>cytoplasm, Collagen-<br>containing extracellular<br>matrix | Cytoplasm,<br>Melanosome |
| ***Mpp6*** | MAGUK p55 subfamily member 6 | 1.95 | 2.52 | Plasma membrane | Membrane, Peripheral<br>membrane |
| **Gbas** | Protein NipSnap homolog 2 | -3.07 | 4.79 | Cytoplasm,<br>Mitochondrion | Mitochondrion outer<br>membrane, Cytoplasm |
| **Slc25a1** | Tricarboxylate transport protein,<br>mitochondrial | -2.75 | 3.68 | Mitochondrion.<br>Mitochondrion inner<br>membrane | Mitochondrion inner<br>membrane |
| **Sorl1** | Sortilin-related receptor | -2.15 | 3.48 | Nuclear envelope lumen | Cell membrane, Single-pass type I membrane<br>protein, Endosome,<br>Secreted, Golgi<br>apparatus membrane,<br>Endoplasmic reticulum<br>membrane, Secretory<br>vesicle membrane |
| - | Ig lambda-1 chain V region | -1.99 | 3.39 | N/A | Extracellular space,<br>Plasma membrane |
| **Sfxn3** | Sideroflexin-3 | -1.38 | 3.08 | Mitochondrion | Mitochondrion<br>membrane |
| **Lmbrd1** | Probable lysosomal cobalamin transporter | -2.57 | 3.04 | Plasma membrane,<br>Lysosome, Clathrin-coated endocytic vesicle | Lysosome membrane |
| **Arl8b** | ADP-ribosylation factor-like protein 8B | -1.24 | 3.04 | Synapse, Axon | Late endosome<br>membrane,<br>Cytoskeleton,<br>Lysosome, Axon |
| **Fundc2** | FUN14 domain-containing protein 2 | -2.48 | 2.93 | N/A | Mitochondrion.<br>Mitochondrion outer<br>membrane |
| **Ckap4** | Cytoskeleton-associated protein 4 | -1.22 | 2.85 | Endoplasmic reticulum | Cytoskeleton,<br>Endoplasmic reticulum<br>membrane, Cell<br>membrane |
| **Mtco1** | Cytochrome c oxidase subunit 1 | -1.33 | 2.80 | Mitochondrion inner<br>membrane | Mitochondrion inner<br>membrane |
| ***Hist1h1b*** | Histone H1.5 | -1.17 | 2.78 | N/A | Nucleus, Chromosome |
| ***Hccs*** | Cytochrome c-type heme lyase | -2.23 | 2.58 | Mitochondrion | Mitochondrion inner membrane |
| ***Mcur1*** | Mitochondrial calcium uniporter regulator 1 | -1.64 | 2.55 | N/A | Mitochondrion inner membrane |
| ***Ctss*** | Cathepsin S | -2.27 | 2.53 | Membrane, Lysosome | Secreted, Lysosome |
| ***Rdh11*** | Retinol dehydrogenase 11 | -1.55 | 2.52 | Photoreceptor inner/outer segment membrane | Endoplasmic reticulum membrane |

**Supplementary Table 7:**
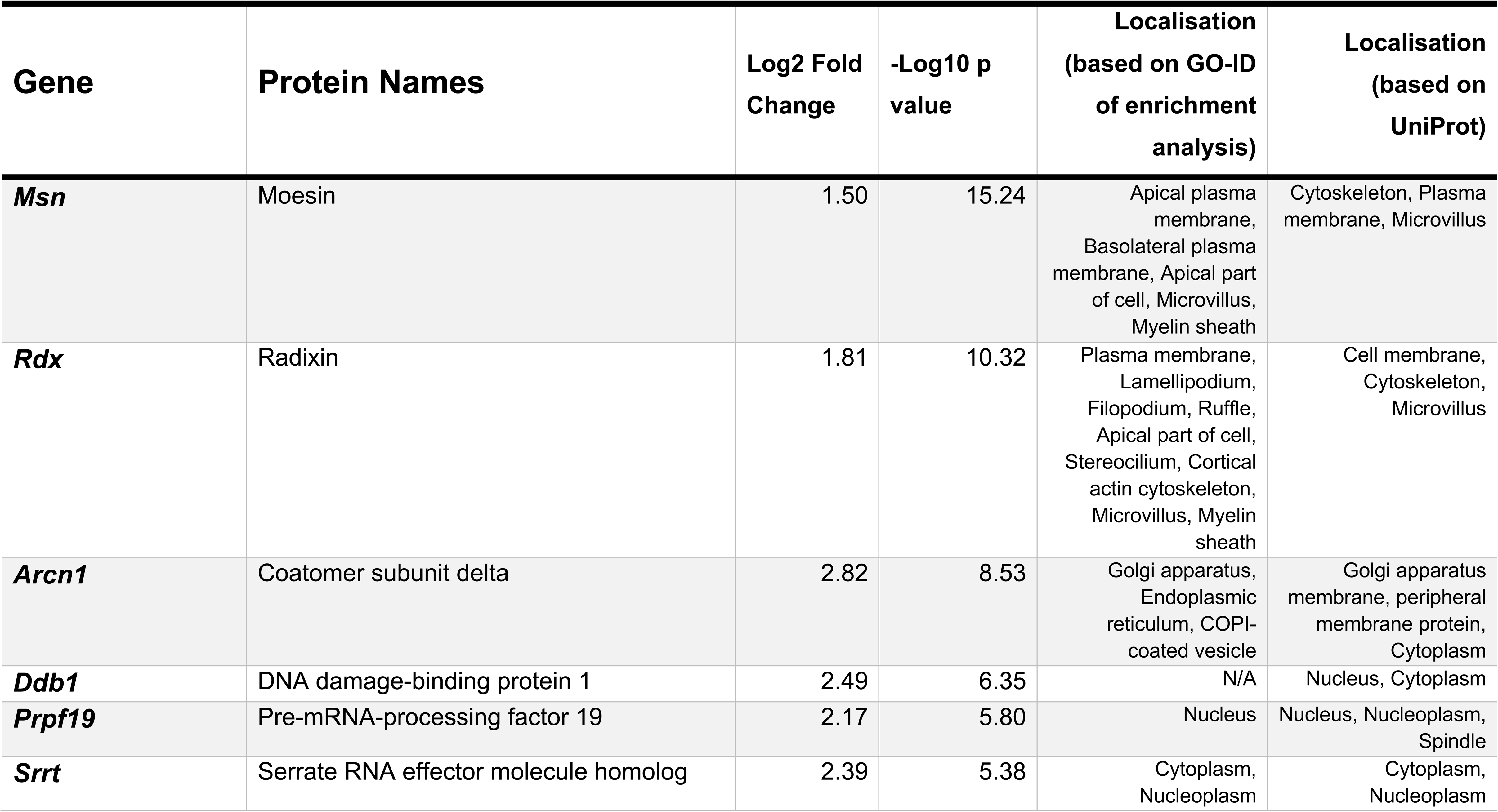
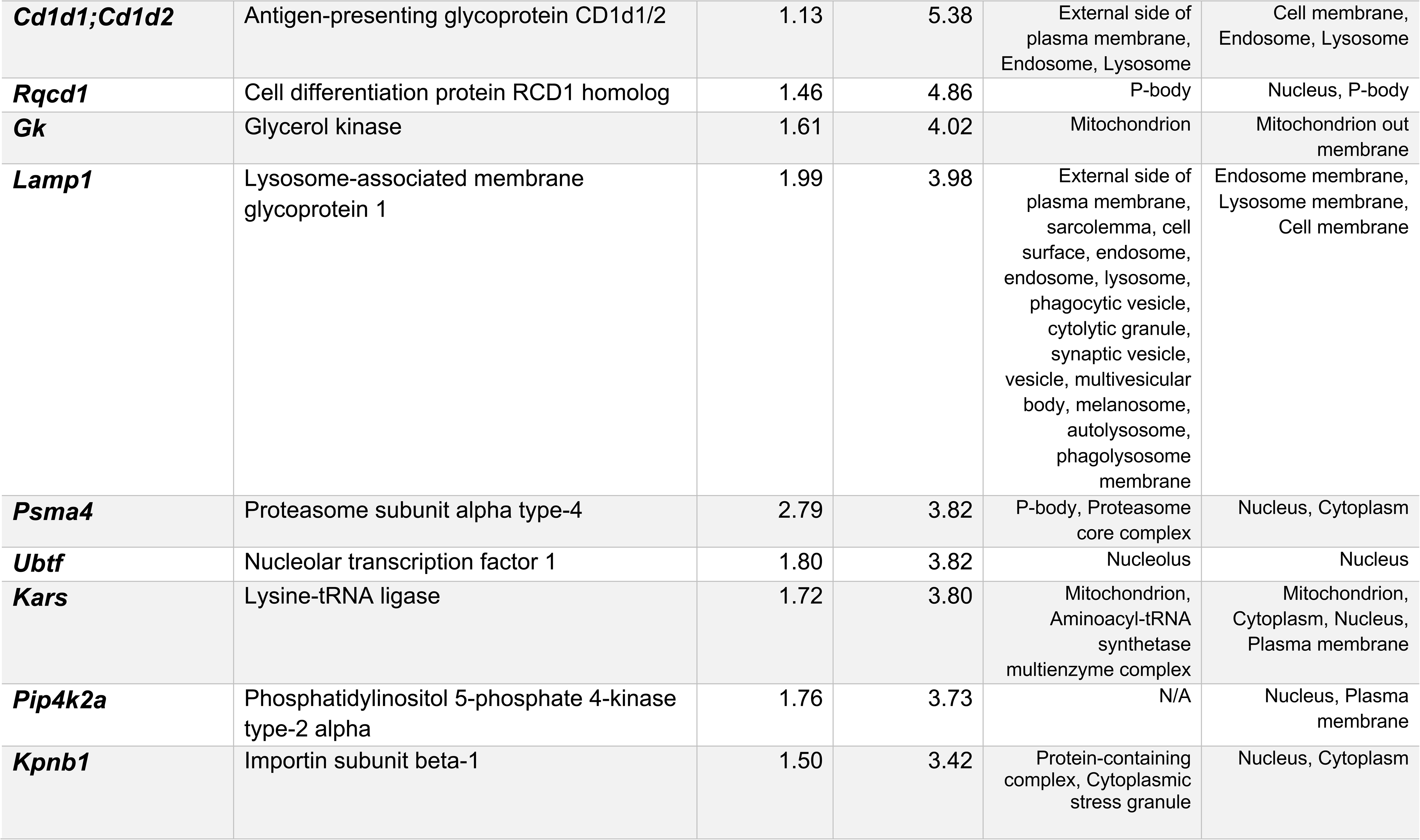
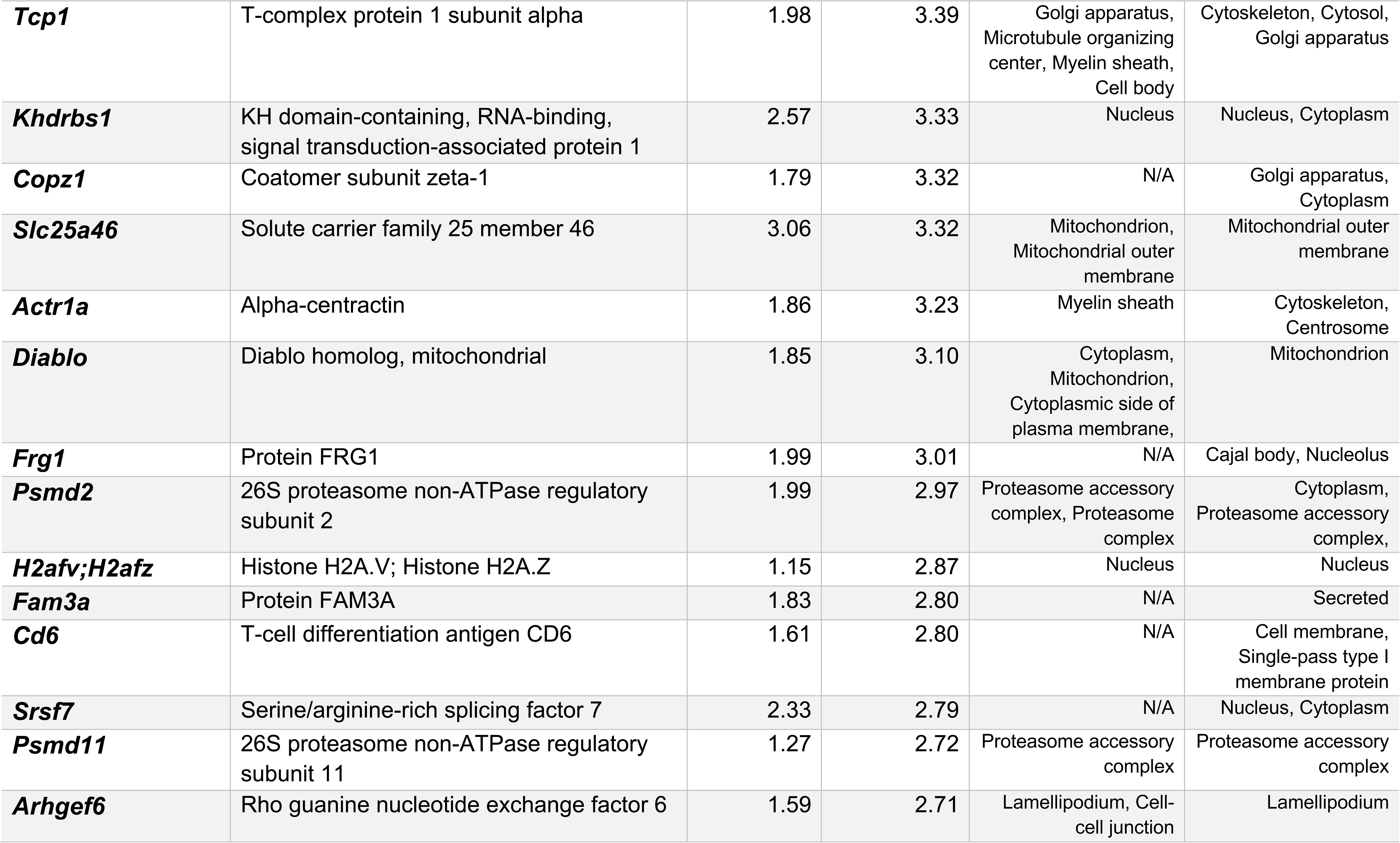
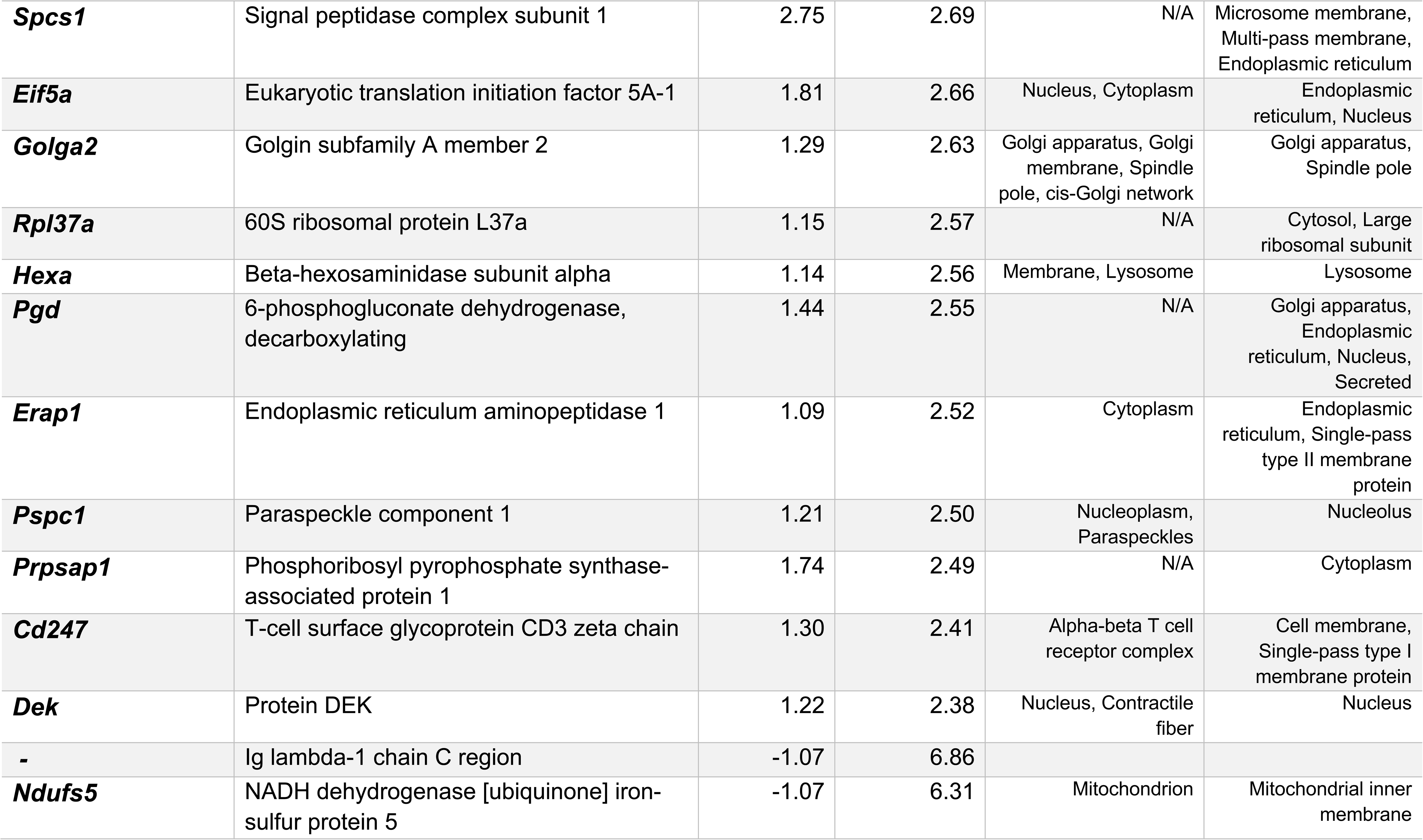
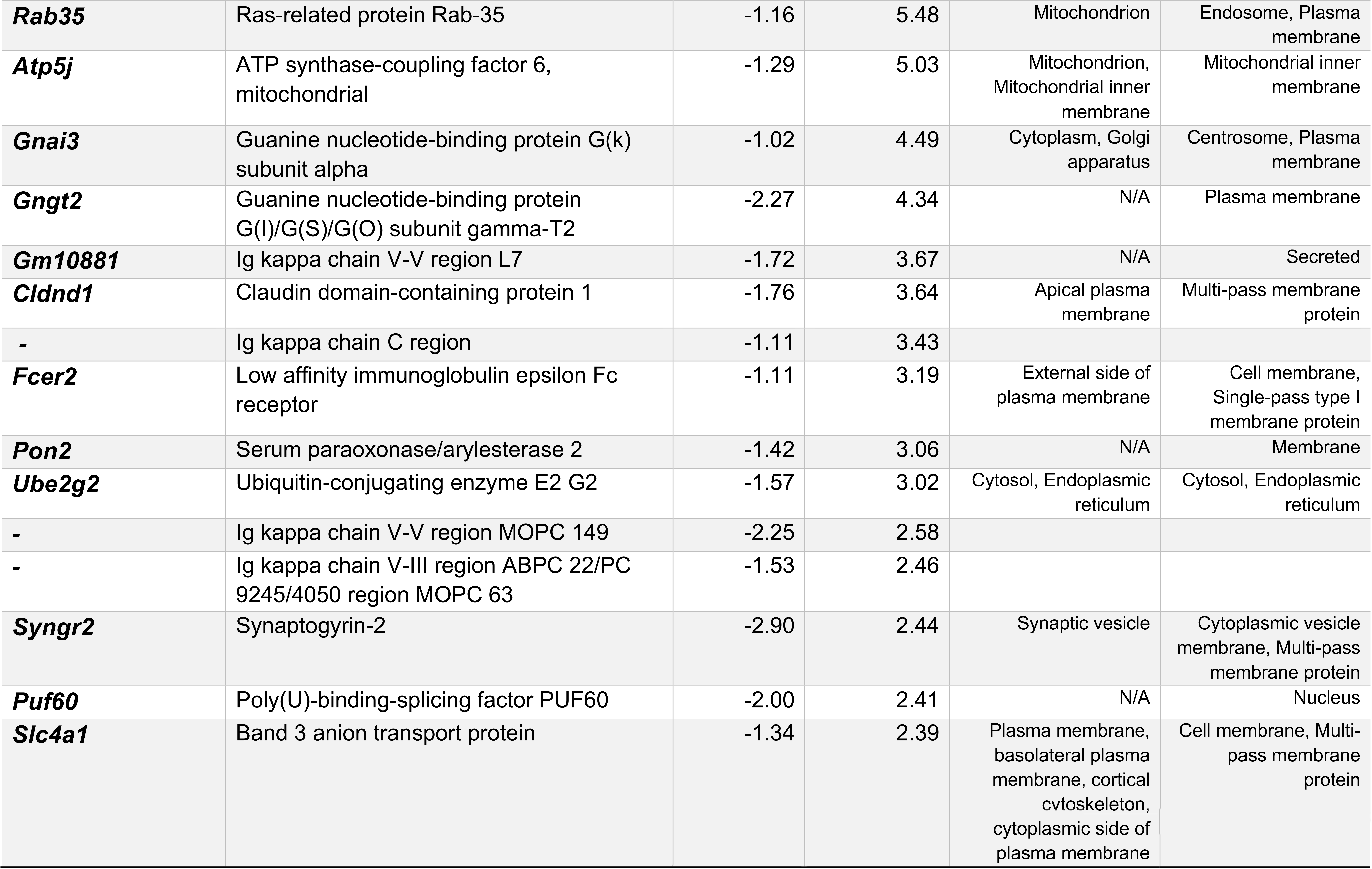
Significantly up/down-regulated proteins in the PM fraction of *Marchf8*^−/−^ B cells.

